# Functional Cell-Type Identification in Neuronal Networks Using High-Density Microelectrode Arrays

**DOI:** 10.64898/2026.04.30.721923

**Authors:** Philipp Hornauer, Lorenzo Davide Dodi, Hsiu-Chuan Lin, Tobias Gänswein, Maria Pascual-Garcia, Manuel Schröter, Andreas Hierlemann

**Author notes:** These authors contributed equally to this work (Philipp Hornauer, Lorenzo Davide Dodi). These authors jointly supervised this work (Manuel Schröter, Andreas Hierlemann).

## Abstract

The reliable identification of neuronal cell types - in particular, the distinction of excitatory (*E*) and inhibitory (*I*) neurons on the basis of extracellular recordings without post-hoc immunostaining or genetic labeling - remains a key challenge in neural-circuit analysis. High-density microelectrode arrays (HD-MEAs) have emerged as a powerful tool to address this issue, enabling simultaneous single-cell and network-level electrophysiology. Here, we present two complementary strategies for establishing cell-type ground truth based on HD-MEA recordings: (i) chemogenetic interneuron activation to label putative inhibitory neurons according to their functional response, and (ii) controlled mixing of excitatory and inhibitory hiPSC-derived populations at defined ratios. A classifier combining action potential waveform morphology and autocorrelogram-based discharge dynamics achieves robust cell-type discrimination in *in vitro* recordings of rat primary cortical cultures and hiPSC-derived networks, as well as in *in vivo* recordings of rat and mouse - i.e., across several species, recording modalities, and preparation types. Applied to unlabeled data, the classifier reveals cell-type-specific network dynamics during bursts, including an inhibition activity signature preceding burst onsets. Leveraging the HD-MEA spatiotemporal resolution, label-free electrophysiological footprint reconstruction enables a morphological characterization of putative *E* and *I* neurons without post-hoc staining. The classification pipeline represents a scalable framework for functional cell-type phenotyping with broad relevance for precision neural-circuit analysis and disease modeling.

## 1. Introduction

Maintaining a balance between excitatory (*E*) and inhibitory (*I*) activity - at the levels of synaptic signaling and cellular composition - is essential for proper neural circuit function, which underlies cognitive processes and adaptive behavior. Imbalances in the *E*/*I* ratio are associated with a range of neurological and neurodevelopmental disorders, including autism spectrum disorder, schizophrenia, and epilepsy (Ben-Ari et al. 2012; Bozzi et al. 2018; Gao and Penzes 2015; Nelson and Valakh 2015). During development, the progressive increase of inhibition drives a transition from early synchronous network activity to the sparser, uncorrelated dynamic characteristics of mature circuits (Chini et al. 2022). A deeper understanding of how alterations of the *E*/*I* balance affect network behavior and function requires tools to modulate interneuron activity and methods to reliably identify the respective cells in the recordings.

Experimental perturbations of the *E*/*I* ratio are commonly achieved pharmacologically by blocking inhibitory synapses, which drives the network toward hypersynchronous, excitation-dominated dynamics (Feinerman et al. 2005; Eytan and Marom 2006). Optogenetics enable more targeted modulation, allowing for selective alteration of GABAergic characteristics both *in vivo* and *in vitro* (Chini et al. 2022; Giraud et al. 2025). Chemogenetics, using Designer Receptors Exclusively Activated by Designer Drugs (DREADDs), offer an alternative with comparable cell-type specificity but simpler experimental implementation, enabling selective activation or inhibition of defined neuronal populations via Gq- or Gi-coupled receptors (Kang et al. 2023; Roth 2016; Scott M. Sternson and Bryan L. Roth 2014).

Another approach to studying defined *E*/*I* ratios involves establishing cultures with precisely controlled neuronal compositions - either through forward programming of inducible neurons with NGN2 or Ascl1 induction (Parodi et al. 2023; Mossink et al. 2022), the use of fluorescence-activated cell sorting approaches (Sukenik et al. 2021; Turko et al. 2019), or the application of dorsal or ventral telencephalic differentiation protocols in monolayer (Crocco et al. 2025) or 3D organoid cultures (Bagley et al. 2017), or the mixing of pure populations(Strong et al. 2023). These strategies are particularly relevant for hiPSC-derived models, where guided differentiation can produce homogeneous excitatory or inhibitory populations that can be combined at defined ratios (Volpato et al. 2018; Hulme et al. 2022).

High-density microelectrode arrays (HD-MEAs) represent a promising tool for studying *E*/*I* interactions, as their large number of tightly spaced electrodes allows for simultaneous analysis of single-cell and network-level activity (Schröter et al. 2025; Obien et al. 2015). Additionally, they enable longitudinal tracking of cultures, facilitating to study developmental dynamics and self-organization principles. However, while electrophysiological methods like patch-clamp enable to determine cell identity through morphology and intrinsic parameters, the analysis of HD-MEA recordings is typically restricted to spontaneous extracellular activity, while neuronal subtypes remain undetermined.

Inferring cell types from extracellular recordings alone has, therefore, become an active area of research. Methods relying on action potential waveform features alone, such as trough-to-peak duration, have shown limited identification performance, particularly for *in vitro* preparations (Becchetti et al. 2012; K. Weir et al. 2015). A growing body of evidence, however, indicates that approaches combining waveform features with spiking dynamics, such as autocorrelograms (ACGs), offer substantially improved discrimination performance (Gonzalez-Ferrer et al. 2025; H. Yu et al. 2025; Lee et al. 2021, 2026). Several groups have demonstrated successful classification for *in vivo* datasets using deep learning or unsupervised methods trained on optogenetically tagged ground-truth libraries, including contrastive learning of jointly embedded waveform and ACG features (NEMO) (H. Yu et al. 2025), self-supervised pretraining with supervised fine-tuning (HIPPIE) (Gonzalez-Ferrer et al. 2025), unsupervised dimensionality reduction across feature types (PhysMAP) (Lee et al. 2026), deep learning for cerebellar cell types (Beau et al. 2025), and vision-language models as few-shot classifiers (Marin-Llobet et al. 2025). SpikeMAP (Giraud et al. 2025) recently introduced a pipeline for cell-type inference in *ex vivo* rodent cortical slice preparations on HD-MEAs.

Overall, most currently available classifiers were developed and validated primarily for *in vivo* settings or slice preparations, and the transfer to *in vitro* preparations of dissociated cells or to other recording scenarios has remained challenging (Becchetti et al. 2012; K. Weir et al. 2015; Beau et al. 2025). HD-MEAs offer unique advantages to address this issue.

Their dense electrode spacing, typically less than 20 µm electrode pitch, improves spike sorting quality/yield and allows for capturing physiologically relevant details, such as the axon initial segment (AIS) position and electrical signal, which substantially enhances cross-unit feature comparability (Bakkum et al. 2019). Moreover, the dense sampling enables to capture axonal signal propagation across the full electrophysiological footprint of individual neurons - a feature inaccessible to standard MEAs or oblong Neuropixels probes. To date, there is no single analysis framework that can be generalized across different preparations and recording scenarios including, for example, dissociated rodent *in vitro* cultures, rodent *in vivo* recordings, and hiPSC-derived networks.

Here, we present two strategies to generate cell-type ground truth for neuronal signals recorded with HD-MEAs: (i) the chemogenetic activation of GABAergic interneurons to assign functional labels in heterogeneous rodent primary cultures, and (ii) the controlled mixing of separately differentiated, excitatory and inhibitory hiPSC-derived populations of defined identity at known ratios. Using the obtained datasets, we created a classification workflow integrating waveform and ACG features that can be transferred across recording scenarios and species, when the corresponding training data are available. Label-free morphological cell-type analysis was demonstrated through electrophysiological footprint reconstruction, and the classifier was applied to reveal cell-type-specific network dynamics in unlabeled cultures and tissues. The complete classification pipeline was implemented as an extension to the publicly available DeePhys toolbox (Hornauer et al. 2024).

## 2. Results

### 2.1. Overview of Cell-type Classification Workflows

Figure 1 depicts the two complementary workflows for assigning excitatory (*E*) and inhibitory (*I*) cell-type labels to neurons recorded with HD-MEAs, each of which is suited for a distinct experimental scenario (Figure 1). The first approach aims at facilitating the labelling of neurons in heterogeneous cultures, such as primary rodent cell preparations, where *E* and *I* neurons co-exist, but cannot be separated *a priori*. The second approach applies to cultures composed of defined, more homogeneous populations of known identity, derived, e.g., through forward programming of human iPSCs (Bocchi et al. 2022). These cellular modelling systems have gained popularity owing to their shorter differentiation time, high reproducibility, and better control over cell-type composition (Zhang et al. 2013; Hulme et al. 2022).

**FIGURE 1.**
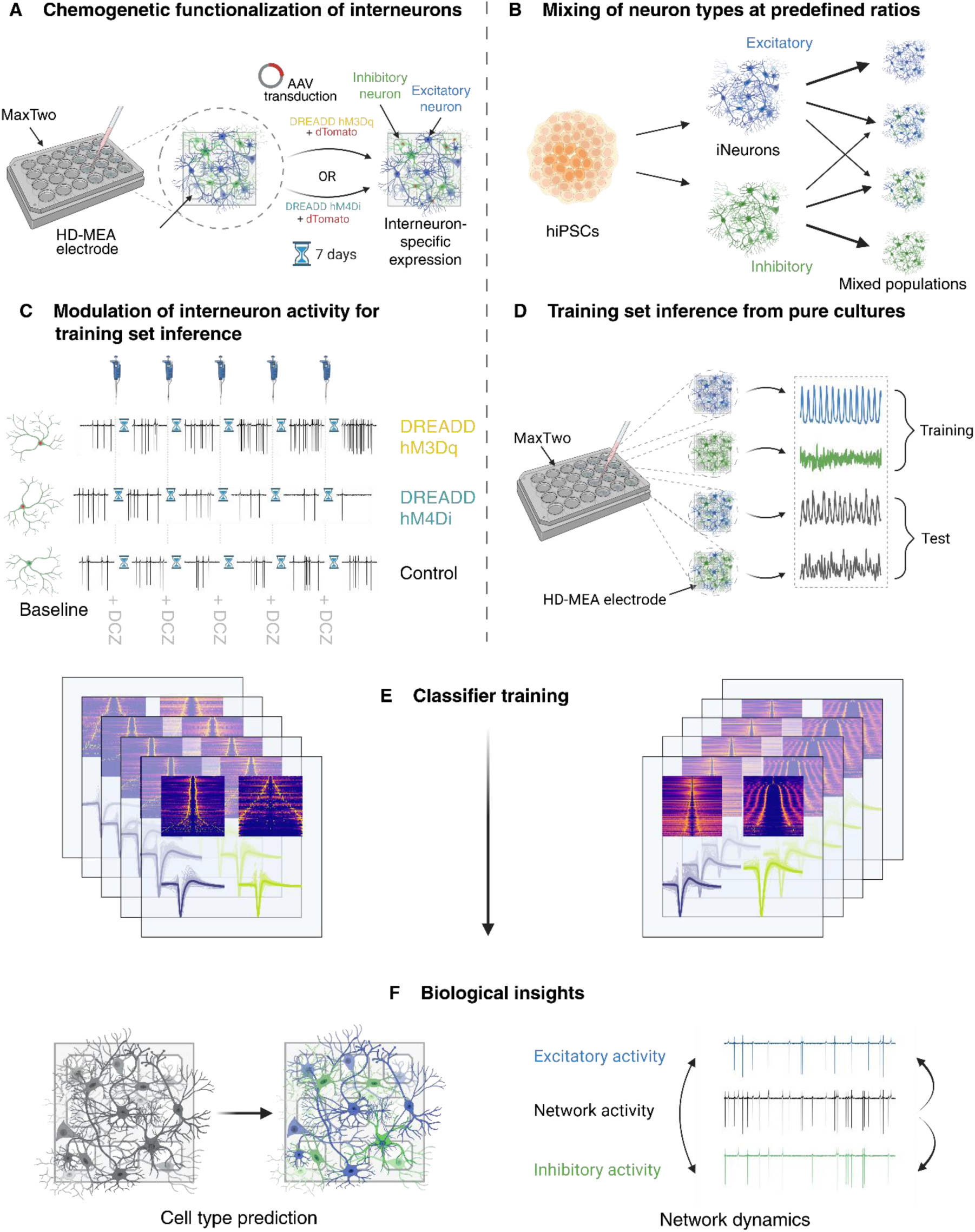
Overview of the two cell-type classification workflows for HD-MEA recordings. **(A)** Heterogeneous neuronal cultures are plated on HD-MEAs (Maxwell Biosystems MaxTwo) and transduced with AAV constructs conferring interneuron-specific expression of either the activatory DREADD hM3Dq or the inhibitory DREADD hM4Di, together with a dTomato fluorescent reporter. **(B)** In the second workflow, excitatory neurons and inhibitory neurons (iNeurons) are derived separately from hiPSCs via guided differentiation and combined at defined ratios prior to seeding. **(C)** Dose-response recordings under sequential DCZ application capture different FR modulations of DREADD-expressing neurons (hM3Dq: increase; hM4Di: decrease; Control: no change), providing putative cell type labels. **(D)** Mixed-ratio hiPSC-derived cultures are recorded on the MaxTwo platform; pure single-type cultures serve as training data and mixed cultures as test data. **(E)** Ground-truth labels are used to train a classifier on putative excitatory and inhibitory neurons, enabling prediction of cell-type-specific network dynamics based on unlabeled recordings **(F)**.

Cell-type-selective chemogenetics were used to generate putative ground-truth labels in heterogeneous cultures across development (DIV 9-23; Figure 1A, 2A). Neuronal cultures were transduced with adeno-associated virus (AAV) constructs driving interneuron-specific expression of either the excitatory DREADD hM3Dq or the inhibitory DREADD hM4Di, co-expressed with a dTomato reporter **(Figure S1)**. Dose-response recordings were then performed by subsequently applying the chemogenetic actuator deschloroclozapine (DCZ). Neurons exhibiting a significant increase in firing rate (FR) in response to hM3Dq activation were classified as putative inhibitory neurons. This chemogenetic labelling strategy helps to establish a data-driven ground truth, which is based on functional response profiles rather than on morphological analyses, optogenetic activation, or post-hoc immunocytochemical staining.

**FIGURE 2.**
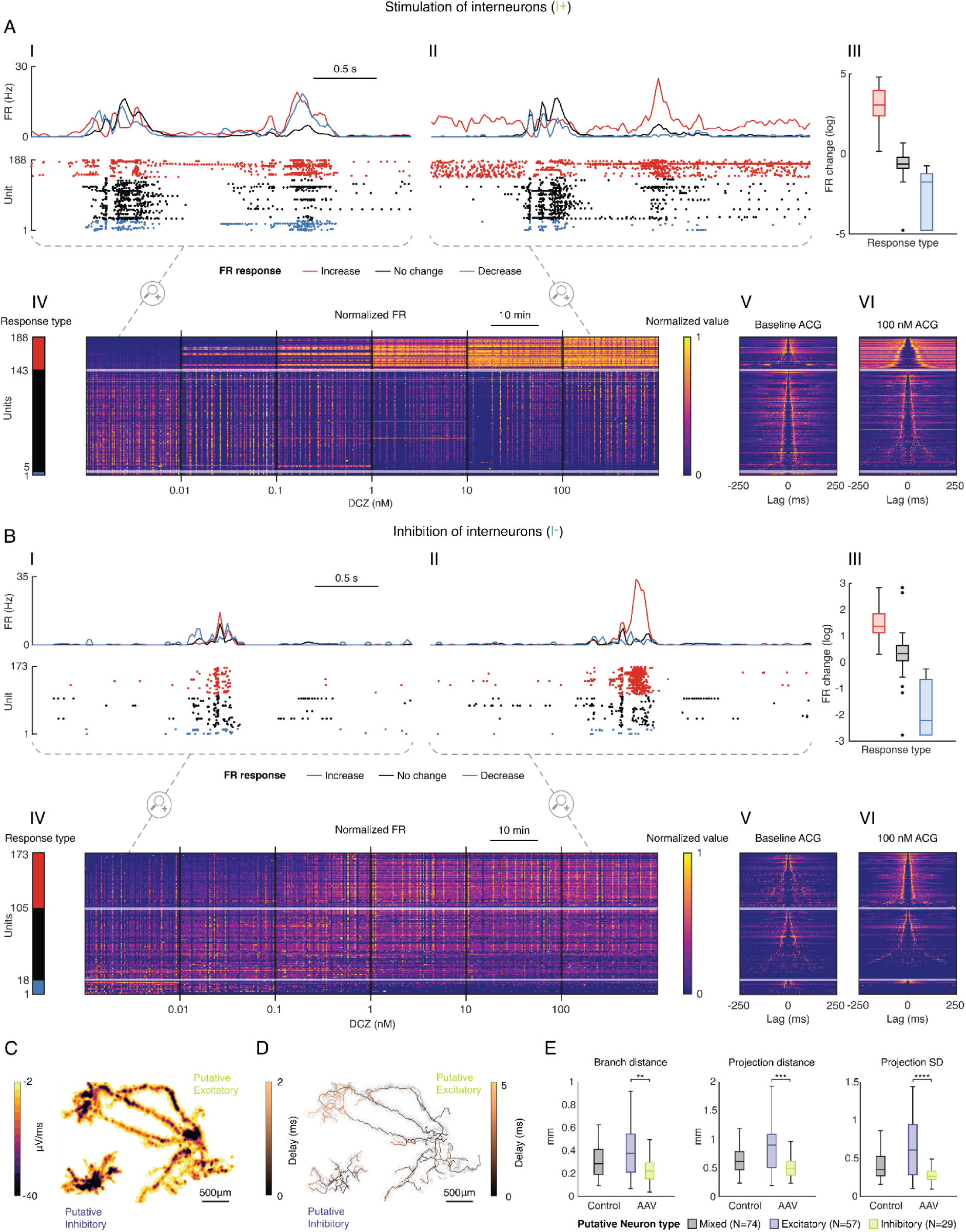
Modulating interneuron activity affects cortical cultures in a dose-dependent manner. Panels **(A)** and **(B)** depict the effects of stimulating (*I*+) or inhibiting (*I*−) the interneuron subpopulation in two example cortical cultures at day *in vitro* 9. For visualization purposes, units were sorted according to their maximum firing rate (FR) response and broadly classified into ‘Increase’ (red), ‘No change’ (black), and ‘Decrease’ (blue). **(I)** Representative network burst in the baseline recording. **(II)** Representative network burst recorded at the highest drug concentration (100 nм DCZ). **(III)** Box plots summarizing the FR changes (log) of the three response groups. FR change was computed as the ratio of post- to pre-stimulation median firing rates. For visualization on a log scale, a pseudocount (half the smallest observed non-zero ratio) was added to all values to accommodate silenced units. **(IV)** Heat map depicting the normalized FRs of all units across the full dose-response experiment (15 s bins). **(V)** Auto-correlograms (ACGs) of the baseline recording. **(VI)** ACGs at the highest drug concentration (bin size: 1 ms, max lag: 250 ms). **(C)** Electrophysiological footprints of representative putative excitatory and inhibitory units, with color map indicating signal amplitude. **(D)** Corresponding skeletonization to the representative units, with color map indicating the delay after action potential initiation (ms). **(E)** Morphological features derived from electrophysiological footprint reconstruction: branch distance, projection distance, and projection standard deviation, comparing Control and AAV-transduced cultures, and between AAV-transduced cultures (** p < 0.01,*** p < 0.001,**** p < 0.0001).

In the second workflow, pure excitatory and inhibitory neuron populations were derived separately via guided hiPSC differentiation and forward programming (Lin et al. 2025) and subsequently mixed at defined E/I ratios before seeding the cells on the HD-MEA surface (Figure 1B, 2B). Recordings of pure cultures (100% glutamatergic or 100% GABAergic) provided a training dataset, while mixed cultures with defined cell ratios were used as test sets to evaluate classifier performance under controlled conditions.

For both classification approaches, HD-MEA recordings were spike sorted using Kilosort 2.5 (Pachitariu et al. 2024), quality controlled using Bombcell (Fabre et al. 2023) and subsequently transferred to DeePhys (Hornauer et al. 2024) for feature inference. Ground-truth labels were then used to train a cell-type classifier. As input features, we used the full action potential waveform shape and the binned ACG. Lee et al. (Lee et al. 2021) demonstrated that UMAP embedding of full waveform shapes outperforms classification based on derived features, such as trough-to-peak duration. Subsequent work has consistently shown that a combination of waveform shape with autocorrelogram-derived temporal dynamics yields superior cell-type discrimination in comparison to using either feature alone (H. Yu et al. 2025; Beau et al. 2025; Gonzalez-Ferrer et al. 2025; Lee et al. 2026). Therefore, we adopted these two features for use in a supervised UMAP framework, which extended the unsupervised approach of WaveMAP (Lee et al. 2021). We concurrently optimized the embedding concerning geometric structure and class separability (Figure 1, row 3). Once trained, the classifier was applied to unlabeled data sets to predict *E* and *I* neuron identity at the single-unit level, enabling downstream analyses of cell-type-specific network dynamics and a label-free morphological cell type characterization through electrophysiological footprint reconstruction.

### 2.2. Generating Cell-Type Labels via Chemogenetic Modulation of Cortical Interneurons

First, we investigated how the chemogenetic modulation of interneuron activity affected cortical cultures at day *in vitro* (DIV) 9 (Figure 2). The hM3Dq DREADD is a Gq-coupled receptor that, upon DCZ binding, depolarizes the expressing cell. Since its expression is driven by the hDlx enhancer, which is selectively active in GABAergic interneurons, only inhibitory neurons are directly activated. Neurons showing a dose-dependent increase in firing rate are, therefore, putative interneurons, whereas those with decreased or unchanged activity respond indirectly to enhanced inhibition. Consistent with this observation, stimulation of interneurons (I+) produced a clear, dose-dependent increase in FR in a distinct neuronal subpopulation, which was identified through a bootstrap permutation test (see Methods for details) comparing each unit’s firing rate at baseline and the two highest DCZ concentrations (Figure 2A).

Comparing bursts at baseline conditions and the highest DCZ concentration (100 nм) in one exemplary cortical culture, revealed that the FR increase in putative interneurons (N=41, mean fold change=40.2) occurred predominantly in the interburst interval, while the activity decrease affected primarily burst-associated firing, highlighting distinct burst phases (Figure 2A I, II). Despite representing a small fraction of recorded units, interneuron activation led to a suppression of the activity of the majority of remaining neurons (N=82/147) in a dose-dependent manner (III, IV). ACG changes were consistent with these FR shifts: putative interneurons were largely silent at baseline conditions, and the characteristic ACG shape could only be observed upon activation (V, VI).

Interneuron inhibition (I−) produced complementary effects, with the most prominent changes occurring during bursts, which became markedly more pronounced upon DCZ addition (Figure 2B, I, II). Overall FR changes, however, were much smaller than after stimulation (III). ACG changes reflected the altered burst structure: units with increased FR developed a sharp ACG peak, consistent with stronger burst-locked firing, while putative interneurons showed reduced autocorrelation (V, VI). Control cultures treated with DCZ showed no systematic changes in activity, ruling out off-target effects of the ligand **(Figures S2 and S3)**. Additionally, we found that dose-response profiles were both more pronounced and more consistent after stimulation of interneurons as opposed to their inhibition (Figure S3).

Leveraging the cell-type labels inferred from the dose-response, we next characterized the functional morphology of putative excitatory and inhibitory neurons via electrophysiological footprint reconstruction (Figure 2C–E, **Figure S4**). AAV-transduced and control cultures showed no significant morphological differences (Kruskal-Wallis, all p > 0.05; N=160 neurons from 12 cultures), ruling out confounding effects through DREADD expression.

Between putative cell types, however, branching distance (0.413 vs 0.253 mm, p=0.004), projection distance (0.946 mm vs 0.592 mm, p=0.0008), and the variability of the individual projection distances of a neuron (projection SD; 0.679 vs 0.334, p<0.0001) differed significantly, while the number of axonal terminals was comparable (18.4 vs 19.4, p=0.979) (Kruskall-Wallis, Dunn’s post-hoc test, Figure S4). These findings demonstrate that HD-MEA-based footprint reconstruction can identify cell-type-specific morphological differences without post-hoc staining, providing independent structural validation of the chemogenetic cell-type inference.

### 2.3. Cell Type Classification is Transferable Across Recording Modalities and Species

Next, we asked whether the chemogenetically inferred labels were sufficient to train a classifier that predicts cell type from action potential waveform and ACG features alone without requiring drug-response data (Figure 3). We first focused on mature cortical rat cultures (DIV 23, N=12 cortical cultures), where FR responses were most robust, and generated training labels from the DCZ dose-response (Figure S3). We found a distinct cluster of units exhibiting a FR increase following DCZ stimulation, which we then considered as putative *I* neurons for the classifier training (Figure 3A, I). As explicit labels for *E* neurons were not available, training data was generated by randomly sampling units outside the *I* cluster. Comparing the waveform (II) and ACG shapes (III) revealed clear differences between the *E* and *I* training data, indicating successful separation of the two cell types. The supervised UMAP embedding of the test data revealed a clear separation of predicted *E* and *I* populations (Figure 3B, I). Predicted *I* neurons displayed narrower action potential waveforms with an earlier, more pronounced repolarization peak (II), consistent with the waveform differences observed in the training data (Figure 3A). ACG shapes also differed markedly between the two populations (III).

**FIGURE 3.**
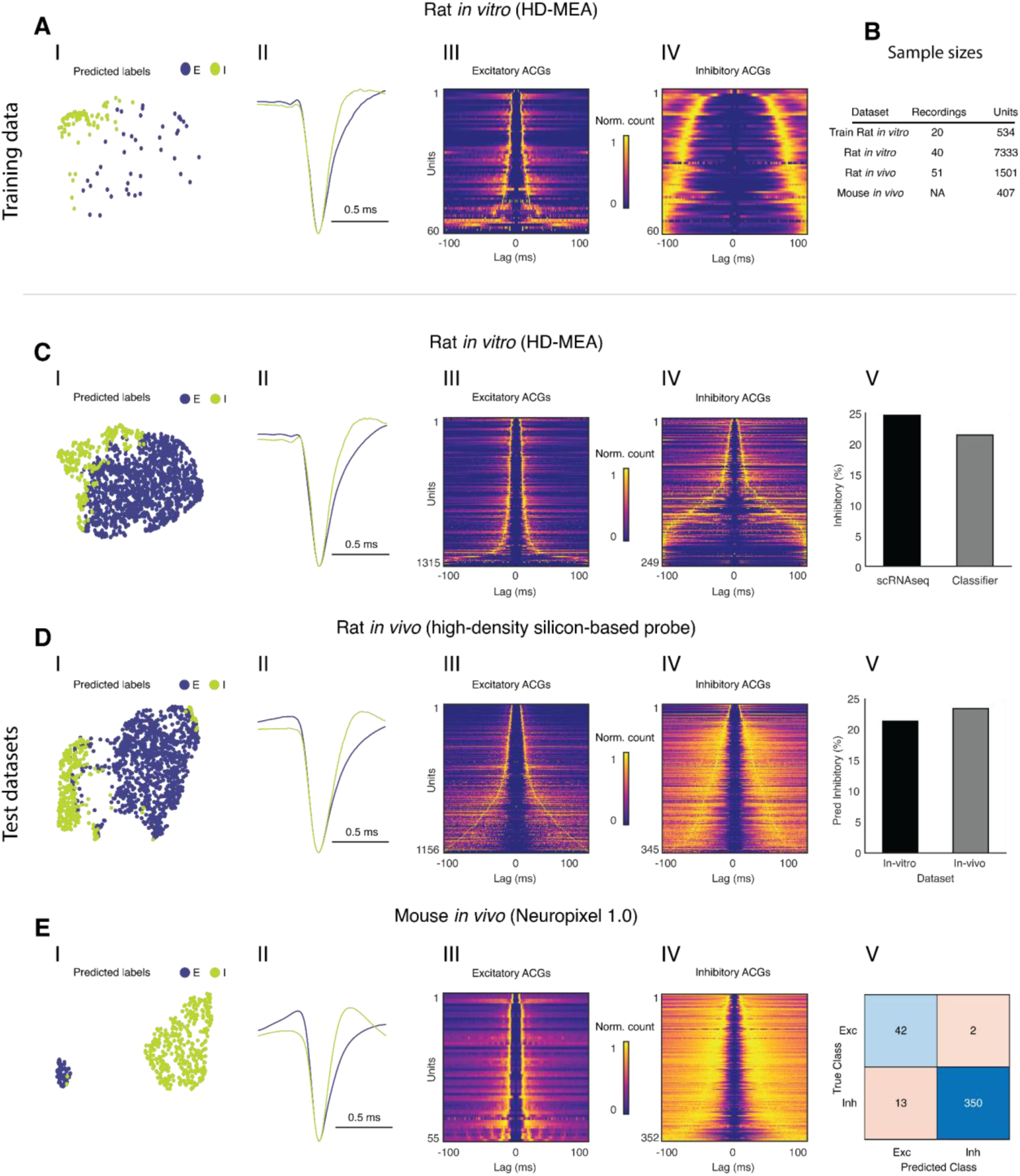
Cell-type classification can be generalized across recording scenarios and species. **(A)** Training labels inferred from chemogenetically stimulated (*I*+) rat cortical cultures (Figure 2A), recorded on HD-MEAs. For clarity, panels A and C show data from a single representative experiment each. **(I)** UMAP embedding of waveform and ACG features, color-coded according to chemogenetic labels (*I*: green, *E*: blue). **(II)** Average normalized waveforms of labeled populations. **(III–IV)** Heat maps of excitatory (III) and inhibitory (IV) ACGs sorted by peak occurrence (bin size: 1 ms, max lag: 100 ms; normalized count). Sample sizes of all data sets are indicated in the accompanying table. **(B)** Table with sample sizes of the utilized data sets indicating recordings and units. **(C–E)** Classifier applied to three held-out test data sets. Each row follows the same layout as (A): UMAP embedding color-coded according to predicted label (*I*), average normalized waveforms (II), excitatory ACG heatmap (III), inhibitory ACG heatmap (IV), and data-set-specific validation (V). **(C)** Rat cortical cultures *in vitro* (HD-MEA). Validation: predicted inhibitory fraction pooled across experiments compared to the proportion derived from scRNA-seq of matched cultures. **(D)** Rat *in vivo* recordings (high-density silicon-based probe). The *in vitro*-trained classifier was applied without retraining. Validation: predicted inhibitory fraction compared between the *in vitro* and *in vivo* datasets. **(E)** Mouse *in vivo* recordings (Neuropixels 1.0). The *in vitro*-trained classifier was applied without retraining. Validation: confusion matrix comparing classifier predictions against optogenetic ground-truth labels.

To validate the classifier predictions quantitatively, we compared the predicted inhibitory fraction across four experiments including two biological batches and two recording time points each (DIVs 9 and 16 and DIVs 16 and 23; 21.4% *I* neurons; **Figure S5**). We found a good agreement between our results and those obtained by single-cell RNA sequencing of matched cultures (24.6% *I* neurons; IV). This agreement suggests that the classifier recovers the expected distribution of *E* and *I* neurons in the tested cortical cultures, though it does not convey any confirmation at the individual-unit level.

Upon applying the classifier - without retraining - to a publicly available rat *in vivo* dataset (Fiáth et al. 2018; Horváth et al. 2021), we found that *E* and *I* neuron populations were clearly separated in UMAP space with more pronounced ACG differences than observed in the *in vitro* rat cortical cultures (Figure 3C, I–III). Although ground-truth labels were unavailable, the predicted inhibitory fraction (23.4%) closely matched the *in vitro* estimate, which suggests transferability and generalizability of the classifier (IV).

Finally, applying the same classifier to a mouse in vivo dataset comprising 407 optogenetically tagged neurons from the CellExplorer database (Petersen et al. 2021), including PV+, SST+, excitatory, axo-axonic PV+, and VIP+ neurons, with preprocessed waveforms and autocorrelograms accessed via the HIPPIE repository (Gonzalez-Ferrer et al. 2025), yielded even more pronounced separation of predicted populations based on their electrophysiological features (Figure 3D, I–III). Optogenetic ground-truth labels allowed for quantitative evaluation: the classifier achieved high precision (350/352 *I* neurons correctly predicted) and high recall (350/363 true *I* neurons identified) for the inhibitory class, with a balanced accuracy of 0.959 and a macro-average F1 score of 0.915 (Figure 3C, IV and **Figure S6**). Importantly, the ground-truth dataset contained only 44 excitatory neurons compared to 363 inhibitory neurons, which inherently limits evaluation of classifier performance for the excitatory class. However, since the classifier was trained to identify inhibitory neurons from the chemogenetic perturbation, with the excitatory class defined by exclusion, the high inhibitory-class performance is the more relevant metric. These results demonstrate that a classifier trained on electrophysiological data from a straightforward chemogenetic perturbation *in vitro* reliably identified interneurons across recording scenarios and species.

### 2.4. Cell-type-specific Activity Dynamics During Network Bursts

Having established putative cell-type labels via chemogenetic modulation of cortical *I* neurons, we investigated the interplay between *I* and *E* subpopulations during network bursts with stimulated interneurons (I+), where FR responses were most consistent (Figure 4). For the purpose of this analysis, all units displaying a significant FR increase across the DCZ dose-response were classified as *I*; all remaining units were treated as *E* neurons. A network-wide I/E activity ratio was derived by grouping the mean FRs of the two subpopulations in 10 ms bins, normalized between 0 and 1. Of note, the I/E rather than E/I formulation was adopted because inhibitory neurons exhibited substantially higher firing rates than excitatory neurons in these cultures, making division by the inhibitory population rate less prone to instability from near-zero denominators.

**FIGURE 4.**
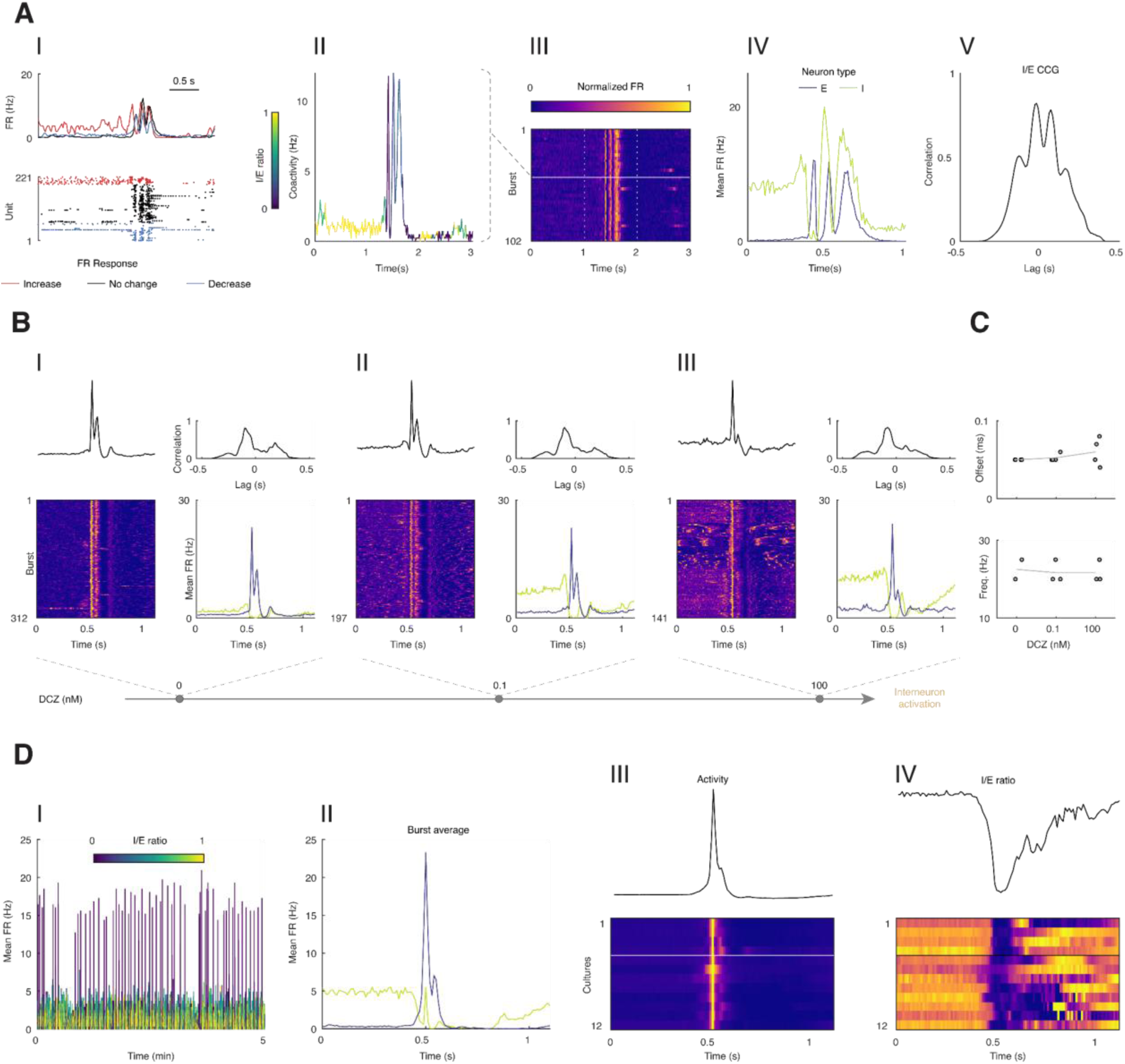
Cell-type-specific activity dynamics during network bursts. **(A)** Analysis pipeline for discrimination of inhibitory (*I*) and excitatory (*E*) subpopulation activity during network bursts. **(I)** Network activity raster color-coded according to FR response type (Increase: red, No change: black, Decrease: blue), with mean FR traces above. **(II)** Close-up of an example burst, highlighting the increase in *I*/*E* ratio (yellow) just before and the sharp decrease (dark blue) during and after the burst. **(III)** Heat map of burst-aligned activity cutouts from a single recording. **(IV)** Mean *I* and *E* neuron FRs inferred from the burst average. ‘Offset’ refers to the time by which *I* activity precedes *E* activity; ‘Period’ refers to the average interval between successive *E* activity peaks during a burst. **(V)** Cross-correlogram (CCG) between *I* and *E* activity derived from (IV). **(B)** Burst dynamics across increasing interneuron activation in cortical cultures at DIV 23 (N=4). Columns **(I)**, **(II)**, and **(III)** correspond to baseline, 0.1 nм DCZ, and 100 nм DCZ recordings, each showing the CCG (top left), burst-aligned activity heat map (bottom left), and mean I and E FR traces (right). **(C)** Summary of offset (top) and firing frequency (1/period, bottom) values across different DCZ concentrations. **(D)** Burst dynamics replicated in unlabeled *in vitro* cultures using predicted cell-type labels. **(I)** Example network activity of a control culture, color-coded by predicted I/E ratio (10 ms bins). **(II)** Predicted *I* and *E* activity traces from the burst average of the same culture. **(III)** Heat map of burst averages across all cultures included in the analysis, with the mean burst characteristics across cultures shown above. **(IV)** Predicted *I*/*E* ratios corresponding to the burst averages in (III). The four cultures at the top were used to generate the *I* training set.

The analysis of rat cortical recordings revealed that burst onsets were consistently preceded by a sharp increase and subsequent drop in the I/E ratio (Figure 4A). Aligning bursts according to cross-correlation and computing the burst average showed that drops in *I* activity did not only mark burst onset but also distinct phases within bursts (III, IV). From these traces we derived two metrics: the temporal offset between the *I* activity peak and the network burst peak, and the period between consecutive *E* activity peaks.

The cross-correlogram (CCG) between *I* and *E* activity captured both the temporal shift and the rhythmic structure of their interaction (V). At baseline conditions, the burst structure was highly consistent in mature cultures (DIV 23), and bursts appeared to be initiated and terminated by spikes in *I* activity, with the *I* subpopulation remaining largely silent during the burst itself (Figure 4B). Increasing DCZ concentration had comparably little effect on burst shape, though at 100 nм, the second burst peak decreased, while the terminal *I* spike intensified. Offset and frequency (20–25 Hz) were consistent across cultures (N=4) and DCZ concentrations (Figure 4C). Notably, younger cultures (DIV 9, **Figure S7**) showed more pronounced burst restructuring with increasing *I* activity, progressing from irregular bursts at baseline through triphasic patterns to ripple-like oscillations at 100 nм DCZ. This suggests that the degree to which interneuron activity shapes burst dynamics depends on the developmental state of the network.

To assess whether these dynamics could be recapitulated using predicted rather than chemogenetically defined labels, we applied the classifier to unlabeled control cultures (Figure 4D). The predicted I/E ratio could be used to reliably distinguish interburst intervals (high I/E) from burst periods (low I/E), and the predicted subpopulation FR traces reproduced the key features identified in ground-truth data: elevated *I* activity during interburst intervals, a sharp drop in *I* activity at burst onset, and brief *I* spikes demarcating burst phases and their termination. Averaging burst activity first within and then across all 12 cultures confirmed that a consistent decrease in the predicted I/E ratio preceded every burst onset, while the activity peak coincided with the I/E minimum - a result that, given the two-level averaging, evidenced both the consistency of the phenomenon and the reliability of the classifier (Figure S7)

### 2.5. Classification of Excitatory and Inhibitory Neurons in Human iPSC-derived Cultures during Maturation

To investigate whether our approach to cell-type classification was applicable to human iPSC-derived cultures, pure excitatory (NGN2-induced glutamatergic) and inhibitory (ASCL1-DLX2-induced GABAergic) neurons were plated as monocultures and mixed cultures (N=30) of defined *E*:*I* ratios (0:100, 25:75, 50:50, 75:25, 100:0) on HD-MEAs and recorded during maturation (Figure 5). At the network level, the *E*:*I* ratio had a pronounced effect on activity patterns: pure excitatory cultures (100:0) displayed robust synchronized bursting, which progressively gave way to lower-amplitude, less structured activity, as the inhibitory fraction increased, with purely inhibitory cultures (0:100) showing largely tonic, non-bursting firing (Figure 5A, DIV 76). Pure culture recordings were used as ground truth for the classifier training (Figure 5B) with mixed cultures held out as test data (Figure 5C).

**FIGURE 5.**
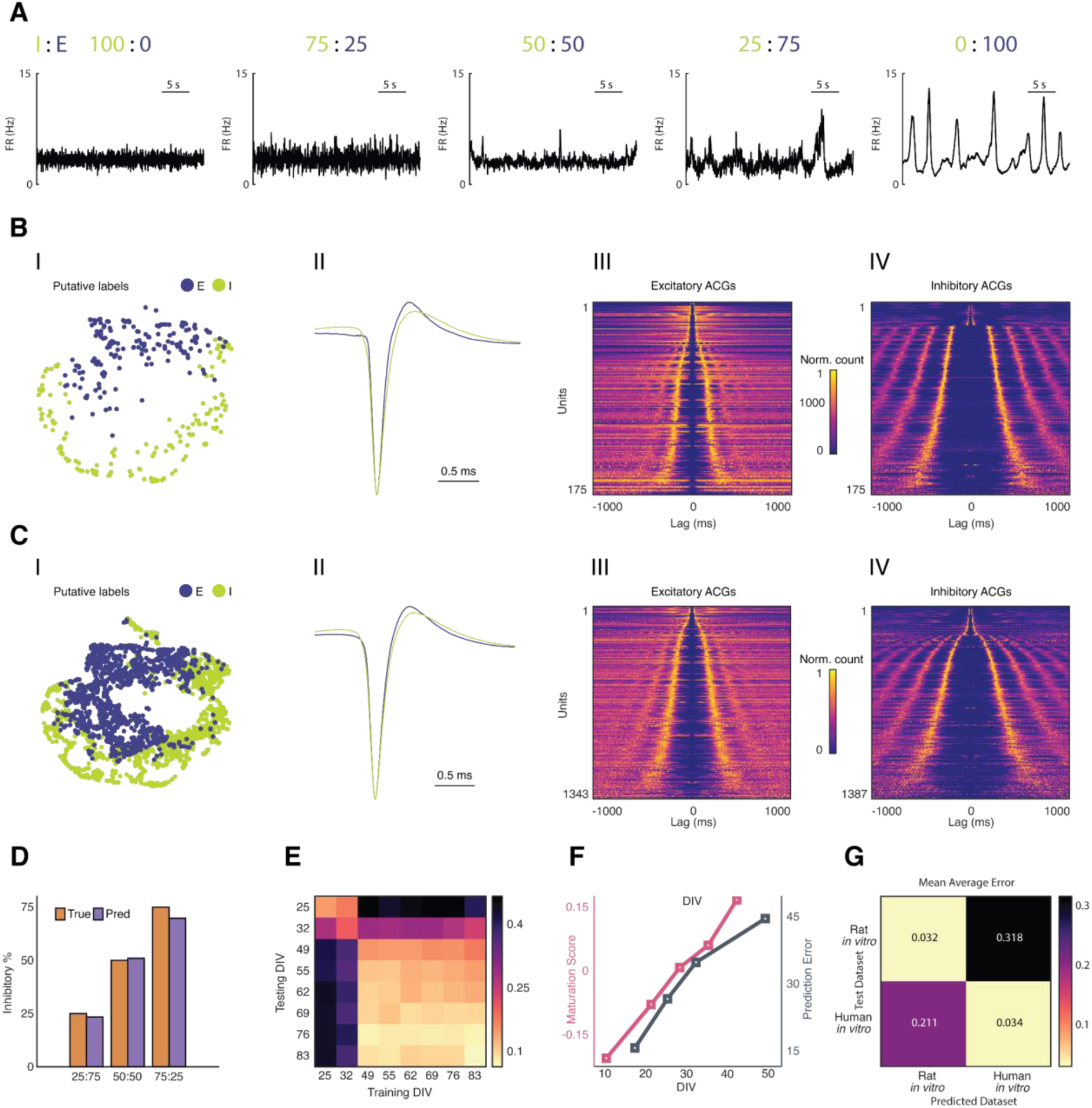
Classification of excitatory and inhibitory neurons in human iPSC-derived cultures during maturation. **(A)** Representative co-activity traces for hiPSC-derived cultures at defined *I*/*E* seeding ratios (100% *I*, 75:25, 50:50, 25:75, 100% *E*) at DIV 76, illustrating the progressive change in bursting dynamics with increasing inhibitory neuron content. **(B)** Ground-truth characterization of pure excitatory and inhibitory hiPSC-derived cultures. **(I)** UMAP embedding color-coded according to cell type. **(II)** Average normalized waveforms, evidencing the broader waveform of inhibitory neurons in comparison to rodent data. **(III–IV)** ACG heat maps for pure excitatory **(III)** and inhibitory **(IV)** cultures (bin size: 2ms, max lag: 1000 ms), with inhibitory neurons displaying prominent oscillatory activity, which was absent in rodent preparations. **(C)** Classifier performance on mixed cultures. **(I)** UMAP embedding of mixed-culture units color-coded according to predicted labels. **(II)** Average normalized waveforms of predicted *E* and *I* populations. **(III–IV)** ACG heatmaps of predicted excitatory **(III)** and inhibitory (IV) units. (D) Predicted fractions of inhibitory cells across mixed cultures compared to true plated I/E ratios at DIV 76. (E) Classification performance during maturation, shown as the mean absolute error per DIV. (F) scRNA-seq maturation score trajectory and prediction error across early development. **(G)** Cross-classification performance matrix displaying the mean average error of each classifier (rat *in vitro* and human *in vitro*) from the expected *E:I* ratio, when applied to both its own and the other species’ dataset.

The UMAP embedding revealed a clear separation of the two populations, and, while waveform differences were present, they were less pronounced than those in primary rat cultures. Notably, iPSC-derived inhibitory neurons displayed broader action potential waveforms than their excitatory counterparts, reversing the narrow-waveform signature that characterizes fast-spiking interneurons in rodent recordings. Comparative scRNA-seq analysis of action potential-shaping ion channels of *E* and *I* neurons in both preparations revealed largely conserved cell-type-specific expression patterns, but also notable exceptions **(Figure S8)**: SCN1A (Na_V_1.1) was strongly enriched in rodent inhibitory neurons but not in hiPSC-derived cultures, while KCNN3 (SK3) showed the opposite pattern.

ACG shapes, however, showed highly consistent differences between cell types across all units: inhibitory neurons displayed prominent oscillatory peaks that were less pronounced or even absent in excitatory neurons. To fully capture these oscillations, ACGs were computed with an extended lag of ±500 ms.

Predicted *E*:*I* ratios in mixed cultures were in good agreement with the known cell plating ratios, which confirmed classifier accuracy (Figure 5D). To assess the effect of maturation on classification, classifiers trained at each recorded DIV were applied to all other time points (Figure 5E). The cross-classification error remained high until DIV 49, which corresponded to the development time trajectory of glutamatergic neurons. Then, it markedly declined, indicating the stabilization of physiological features after this time point. (Figure 5F).

Finally, we assessed the cross-species transferability and generalizability of both classifiers (rat *in vitro* and hiPSC-derived) across the two datasets with validation against independent references: scRNA-seq data were used for rat *in vitro* cultures and plating ratios for hiPSC mixed cultures (Figure 5G). The rat *in vitro* classifier performed well on its own held-out data, giving a mean average error of 0.032, but showed lower performance upon application to human cultures, yielding a mean average error of 0.318. The opposite pattern was observed for the hiPSC-derived classifier, which performed strongly on human mixed cultures, 0.034, but showed lower performance upon application to the rodent preparation, 0.211. Together, these results suggest that electrophysiological feature distributions are both species- and preparation-specific, making matched training data essential for reliable classification of data sets.

### 2.6. Cell-type-specific Oscillatory Dynamics for Different *E*:*I* Ratios

Applying the classifier to mixed hiPSC-derived cultures enabled cell-type-specific characterization of the dynamics observed in the ACGs (Figure 6). Across all *E*:*I* ratios, several distinct ACG patterns were apparent. Most notably, a clearly rhythmic, oscillatory pattern was present in the majority of inhibitory units. The presence of seemingly distinct ACG profiles that were cell-type-specific prompted us to develop an ACG-based subtype classifier to determine whether cell-type identity and network composition would influence the proportions and development of such subtypes. The classifier was designed to assign putative units to one of four classes: irregular, weak oscillatory, oscillatory, and bursty.

**FIGURE 6.**
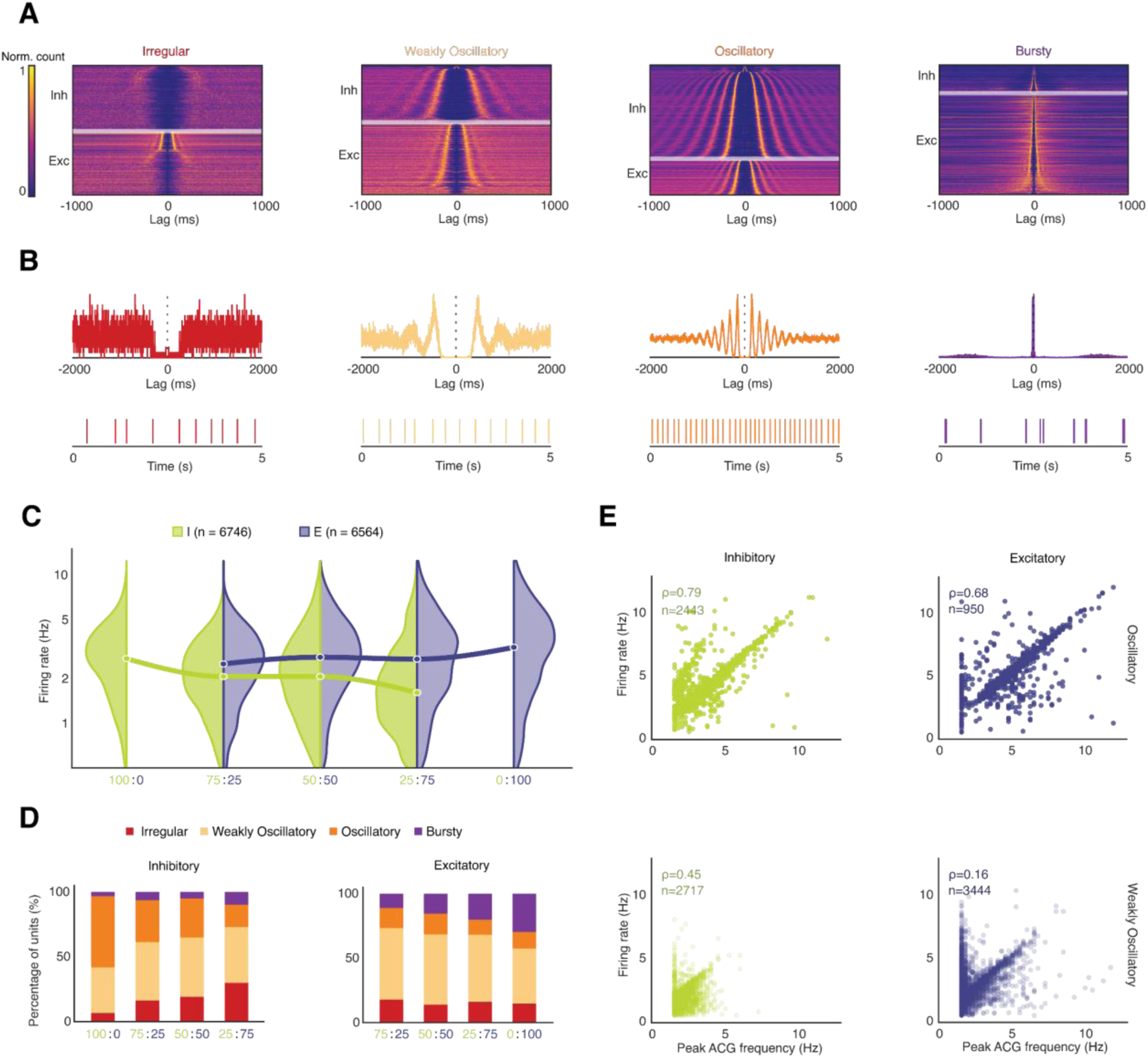
Cell-type-specific oscillatory dynamics across *I*:*E* ratios in hiPSC-derived mixed cultures. **(A)** Heat maps of ACGs of inhibitory (top) and excitatory (bottom) neurons, sorted by peak occurrence (bin size: 2 ms, max lag: 1000 ms; normalized count), assigned to four classes of sub-types: irregular, weak oscillatory, oscillatory and bursty. **(B)** Representative ACGs (top; bin size: 2 ms, max lag: 2000 ms) and spike trains (bottom) of a unit belonging to each of the 4 ACG-based subtypes. **(C)** Split violin plot representing the mean firing rate of units across cultures with different *I*:*E* ratios, separately for inhibitory (left, green, N = 6,746) and excitatory (right, indigo, N = 6,564) cells. Median points of each split violin are connected to the median of the neighbouring split violins of the same cell type. **(D)** Stacked bar plots for each cell type (excitatory left, inhibitory right), showing the distribution of ACG-based sub-types across different *I*:*E* ratio conditions. **(E)** Scatter plots of mean firing rate and peak ACG frequency according to cell type (excitatory, left; inhibitory, right) and oscillatory ACG subtype (weak oscillatory, bottom; strong oscillatory, top).

Upon examining the contribution of each ACG subtype for the different culture conditions, a distinct trend emerged. In purely inhibitory cultures, around 90.1% of units exhibited oscillatory profiles (weak or strong), which became progressively noisier and burstier with the addition of excitatory neurons. Excitatory units underwent an analogous shift in the opposite direction: pure excitatory cultures contained a large proportion of bursty units (29.6%), which gradually gave way to oscillatory subtypes, as inhibitory neurons were introduced. Notably, the proportion of irregularly firing units within the excitatory population did not decrease upon addition of inhibitory cells, remaining near baseline levels (∼15%).

To further validate the rhythmic properties of units classified as oscillatory subtypes, we examined the correlation between peak frequency in the ACG spectrum and mean firing rate in the two subpopulations. Units assigned to the stronger oscillatory subtype exhibited a considerably higher Spearman’s rho (*I*: ⍴=0.79, *E*: ⍴=0.68, vs *I*: ⍴=0.45, *E*: ⍴=0.16) than their weakly oscillatory counterparts, which supported the validity of our classification.

Both cell types maintained relatively stable mean firing rates (∼2-3 Hz) regardless of the network composition and varying ratios of excitatory to inhibitory neurons. Nevertheless, cell-type-specific trends were consistent across *E*:*I* conditions: inhibitory neuron firing rates were highest in purely inhibitory cultures (median of 2.73 Hz) and declined progressively with increasing excitatory proportions, reaching their lowest values for the 75:25 ratio (median of 1.62 Hz). Conversely, excitatory neuron firing rates peaked in purely excitatory cultures (median of 3.26 Hz) and decreased as the proportion of inhibitory neurons grew, reaching their minimum for the 25:75 ratio (median of 2.52 Hz). These findings suggest that the presence of excitatory neurons is associated with reduced inhibitory neuron firing rates across different network compositions.

Firing rate distributions further could be used to distinguish the two predicted populations across all mixture conditions. Predicted inhibitory neurons displayed consistently lower mean firing rates than their excitatory counterparts across all *E*:*I* ratios - a finding that contrasts to the higher firing rates typically observed for rodent inhibitory neurons and that likely reflects intrinsic electrophysiological differences in the ASCL1-DLX2 GABAergic line.

## 3. Discussion

In this study, we developed and validated a pipeline for inferring excitatory and inhibitory neuron identity from extracellular HD-MEA recordings alone, using two complementary ground-truth strategies, We also demonstrated transferability and generalizability across recording scenarios, species, and neuronal preparation nature or cell line origin.

The use of DCZ as a chemogenetic actuator *in vitro* proved effective at concentrations that were several orders of magnitude lower than previously reported for *in vitro* applications (J. S. Weir et al. 2023; Nentwig et al. 2022), with effects already detectable at 0.01 nм. This finding is relevant, as higher concentrations increase the risk of having off-target effects (Nagai et al. 2020). The dose-response profiles revealed that interneurons fire predominantly during interburst intervals under both stimulation and inhibition conditions, which is consistent with PV+ interneurons being the dominant GABAergic subtype in the rodent cortex (Rudy et al. 2011). PV+ interneurons are known to be hyperpolarized at baseline *in vitro* (Tepper et al. 2010) and capable of sustaining very high FRs upon activation (Tepper et al. 2010; McCormick et al. 1985). The low baseline firing rate of interneurons likely contributes to the difficulty of *in vitro* cell-type classification, as the ACG features that distinguish fast-spiking interneurons *in vivo* are less pronounced when these neurons fire only sparsely.

Label-free reconstruction of axonal morphology through HD-MEA-derived electrophysiological footprints revealed that putative interneurons had smaller but more densely branched axonal arbors with shorter projection lengths, consistent with PV+ interneuron morphology (Tepper et al. 2010; Zeng and Sanes 2017; Kepecs and Fishell 2014; Y. Kawaguchi 1993). The absence of significant differences between AAV-transduced and control cultures rules out that there is a major effect of DREADD expression on morphology. Notably, cell-type-specific morphological differences derived from electrophysiological footprints have previously been reported only in pure cultures of known cell type identity (Radivojevic and Rostedt Punga 2023). The results presented here demonstrate that such differences can also be identified in heterogeneous, mixed cultures on the basis of predicted cell-type labels alone. These findings establish HD-MEA-based footprint reconstruction as a means of providing independent structural validation of functional cell-type assignments in intact networks. This capability is unique to 2D high-density array recording platforms and inaccessible to conventional MEAs or shank-type *in vivo* probes.

The systematic characterization of the *I* and *E* subpopulation interplay during network bursts revealed a consistent pattern: *I* activity spikes preceded burst onsets, followed by *I* silence and rhythmic *E* activity peaks, the frequency of which changed with network maturity (∼10 Hz in immature, ∼20 Hz in mature cultures). These findings align well with patch-clamp studies linking GABAergic activity to focal seizure onset (de Curtis and Avoli 2016; Grasse et al. 2013), interneuron-mediated synchronization of pyramidal cells (Cobb et al. 1995), and interleaved *E*/*I* spiking during *in vitro* seizure-like events that are consistent with interneuron depolarization blocks (Jokubas Ziburkus et al. 2006). To the best of our knowledge, our findings are the first systematic network-scale demonstration of this phenomenon with HD-MEAs. Notably, the failure to recapitulate these dynamics in the rat *in vivo* dataset probably reflects a fundamental difference in network organization: *in vivo* recordings sample a sparse, spatially restricted subset of a much larger circuit, precluding the dense population-level sampling required for a meaningful *I*/*E* activity ratio estimate.

Previous attempts to classify *E* and *I* neurons based on MEA recordings yielded inconclusive results (K. Weir et al. 2015; Becchetti et al. 2012), largely because waveform shape alone is insufficient, particularly *in vitro* (Beau et al. 2025). By combining waveform and ACG features in a supervised UMAP framework, and by using chemogenetic perturbations as partial ground truth, we achieved reliable classification that was transferable across batches, maturation stages, and, importantly, could be applied to *in vivo* data sets of two different recording devices, which the classifier was never trained on (Jun et al. 2017; Fiáth et al. 2018). The high precision but moderate recall on the opto-tagged mouse data set (350/352 predicted interneurons correctly identified; 350/363 true interneurons captured) reflects the expected asymmetry of training labels derived exclusively for the inhibitory class, and is consistent with results of comparable approaches (Beau et al. 2025).

Several features distinguish the present approach from concurrent and previously reported methods. First, the strategy to chemogenetically establish a ground-truth enables classifier training solely based on *in vitro* data, which eliminates any dependence on optogenetically tagged *in vivo* data sets that are characteristic for NEMO (H. Yu et al. 2025), HIPPIE (Gonzalez-Ferrer et al. 2025), and the Beau et al. framework (Beau et al. 2025). This fact is significant for two reasons: (i) we demonstrated that establishing a classification method in dissociated cultures, where electrophysiological differences between cell types are less pronounced (cf. UMAP separations in Figure 3), also yields a viable solution for *in vivo* classification; (ii) establishing a cell-type ground truth in dissociated *in vitro* preparations through optogenetic tagging entails considerable experimental complexity that has so far limited the efforts in this area. Second, the developed classifier could be applied across three distinct recording platforms at different sampling rates (HD-MEA at 10 kHz, rat *in vivo* probe at 20 kHz, Neuropixels at 30 kHz), which demonstrates the robustness to technical variability, while domain-adaptation strategies, employed, e.g., by HIPPIE, were not needed. Third, in contrast to the deep learning architectures used by NEMO, HIPPIE, and Beau et al., the UMAP-based approach is computationally efficient and transparent, requiring minimal training data and hardware resources. While deep learning approaches may achieve higher absolute accuracy on large, well-annotated *in vivo* datasets, the framework presented here is better suited for the data-limited regime, which is typical for *in vitro* experiments. Notably, the analyses presented here were performed on automatically curated rather than hand-picked units, which renders the classification task considerably more challenging than in studies using manually selected high-quality recordings. However, the scenario of our study also more realistically reproduces general experimental conditions. Finally, compared to SpikeMAP, which was developed based on data of acute cortical slices with preserved cytoarchitecture, the present work addresses the more challenging scenario of dissociated neuronal cultures and additionally extends to human iPSC-derived preparations.

The broader action potential waveform of hiPSC-derived inhibitory neurons and their lower firing rates compared to their excitatory counterparts - both contrasting with the narrow waveforms and high firing rates that characterize rodent fast-spiking interneurons - may be partly explained by different ion channel expression. Our scRNA-seq data revealed strong differences in the expression of SCN1A (encoding NaV1.1) and KCNN3 (encoding the SK3 calcium-activated potassium channel) between the two populations. *In vivo*, Na_V_1.1 is preferentially expressed in GABAergic interneurons, particularly PV+ and SST+ subtypes, where it is critical for action potential initiation and sustained high-frequency firing. A loss of NaV1.1 function selectively impairs interneuron excitability and reduces peak firing rates (F. ^H^. Yu et al. 2006; Ogiwara et al. 2007; Dutton et al. 2013^)^. Reduced or immature SCN1A expression in ASCL1-DLX2-derived neurons could, therefore, contribute to both the broader action potential waveforms and the lower firing rates observed here. KCNN3 encodes an SK channel that mediates the calcium-dependent after-hyperpolarization and is essential for maintaining firing regularity and pacemaker precision in neurons that express it (Wolfart et al. 2001). Different KCNN3 expression in the two populations may contribute to the prominent oscillatory ACG profiles observed in the inhibitory population. Together, the observed channel expression differences provide a molecular basis for the electrophysiological features that distinguish the two cell types in our hiPSC-derived system. These differences may also explain why classifiers trained on rodent data, where interneurons express these channels at levels sufficient for the canonical fast-spiking phenotype, cannot be directly transferred to human iPSC-derived preparations.

The observation that both cell types maintained relatively stable mean firing rates (∼2–3 Hz) across different *E*:*I* conditions - despite the decrease of cell-type-specific firing rates with increasing hetero-type proportion - is consistent with homeostatic regulation of network activity in hiPSC-derived cultures (Sukenik et al. 2021; Sasaki et al. 2019). Whether the rate reductions reflect network-level mechanisms analogous to those described in inhibition-stabilized networks (Tsodyks et al. 1997), or are due to alternative factors, such as competition for trophic support or differences in synaptic drive across seeding compositions, remains to be determined.

Several limitations of the present work should be acknowledged. The chemogenetic ground truth, while validated against optogenetic tagging, only provides labels for the inhibitory class; excitatory labels are inferred indirectly by exclusion, meaning that the two classes have asymmetric ground-truth quality. Future work could employ dual labeling strategies to obtain positive labels for both populations. The classifier currently only distinguishes broad excitatory and inhibitory categories; extension to finer-grained subtypes (e.g., PV+, SST+, VIP+ interneurons) would require strategies that enable establishing subtype-specific ground-truth. The lacking cross-species transferability of the classifier, while informative, limits applicability and motivates the development of domain-adaptation approaches or universal feature spaces that are invariant to species and neuronal preparations.

Looking forward, the framework established here is well positioned for several translational applications. Disease models, based on hiPSC-derived neurons, particularly those modeling *E*/*I* imbalance in autism spectrum disorder, schizophrenia, or epilepsy, would benefit directly from the ability to track cell-type-specific dynamics across development and upon pharmacological perturbation. Integration with high-density *in vivo* probes could extend the approach to chronic recordings in behaving animals, further bridging the gap between *in vitro* phenotyping and *in vivo* circuit function.

## 4. Experimental Section/Methods

### Primary Rat Cortical Cultures

All animal procedures to harvest rodent primary neurons were approved by the veterinary office of the Canton Basel-Stadt under Swiss federal animal welfare laws. HD-MEAs were sterilized in 70% ethanol (30 min), washed with deionized (DI) water, coated with 0.05% poly(ethyleneimine) in borate buffer (pH 8.5, 1 h), and subsequently with 0.02 mg/mL laminin in Neurobasal medium (30 min, 37 °C). Cortices of E18 Wistar rat embryos were dissected in ice-cold HBSS, dissociated with 0.25% trypsin-EDTA, and plated at 24,000 cells per array. After 1 h, 1 mL plating medium was added (Neurobasal/horse serum/GlutaMAX/B-27 Plus, 45:5:0.125:1). Twice a week half the volume of the medium was exchanged using BrainPhys-based growth medium (BrainPhys/SM1/Penicillin-Streptomycin, 97:2:1). Cultures were maintained at 37 °C, 5% CO₂.

### Human iPSC-Derived Neuron Cultures

Excitatory neurons were derived from the NGN2-inducible line 409B2-iNGN2 (provided by B. Treutlein, ETH D-BSSE; (Lin et al. 2025)) and inhibitory neurons from the ASCL1-DLX2 GABAergic line iLB-C-133bm-s4 (provided by O. Brüstle, Universitätsklinikum Bonn; (Peitz et al. 2020)). HD-MEA plates were coated with poly-L-ornithine (PLO), followed by laminin (1:500, L2020, Roche, overnight). NGN2 neurons were seeded at 10,000 cells/cm², induced with doxycycline (4 µg/mL) until DIV3, and carried through a defined medium progression from DMEM/F12-based induction medium (DIV 0–2) to Neurobasal/B-27/BDNF/NT3 maintenance medium (DIV 3+), with Ara-C (2 µM) added until DIV 5 and FBS (2.5%) from DIV 9. GABAergic neurons were seeded at 20,000 cells/cm², induced with doxycycline (4 µg/mL) until DIV7, and carried through a defined medium progression from DMEM/F12-based induction medium (DIV 0–2) to Neurobasal/B-27/BDNF/NT3/Lam maintenance medium (DIV 3+), with Ara-C (2 µM) added until DIV 5, ROCK inhibitor until DIV3, DAPT from DIV3 until DIV7, and FBS (2.5%) from DIV 9. On DIV 7, cells were re-seeded in co-cultures with primary rat astrocytes (1:2 ratio) at the specified glutamatergic:GABAergic ratios (100:0, 75:25, 50:50, 25:75, 0:100). Samples sizes were N=6, N=6, N=9, N=6, N=3, respectively. Weekly recordings were performed from DIVs 10 - 83.

### Adenovirus-associated-virus (AAV)-mediated Transduction of Chemogenetic Constructs

To enable the selective activation of GABAergic interneurons, we used an AAV vector that induced the expression of the designer receptors only upon activation by designer drugs (DREADD) hM3Dq under the transcriptional control of the human distal-less homeobox (hDlx) enhancer element and the human beta-Globin (HBB) minimal promoter fragment (ssAAV-1/2-hDlx-HBB-chI-HA_hM3D(Gq)_2A_NLS_dTomato-WPRE-SV40p(A), serotype 1; Addgene #83897; Viral Vector Facility Zurich). The AAV vector additionally induced the expression of the fluorescent protein dTomato containing a nuclear localization signal, which allowed us to identify successfully transduced neurons. To selectively silence GABAergic interneurons, we induced the expression of the DREADD hM4Di through an otherwise identical AAV vector (ssAAV-1/2-hDlx-HBB-chI-hM4D(Gi)_2A_NLS_dTomato-WPRE-SV40p(A), serotype 1; Addgene #83896; Viral Vector Facility Zurich). We transduced cortical cultures one week before the first electrophysiological recording to ensure reliable expression of DREADDs. The medium was removed entirely, and 0.5 mL of growth medium containing the respective AAV at MOIs of 2.3×10^6^ (hM3Dq) or 3.1×10^6^ (hM4Di) was added. After an incubation period of 4 hours at 37 °C, another 0.5 mL of growth medium was added. A full medium exchange was performed 3 days later to remove the remaining AAVs from the culture.

### High-density Microelectrode Array (HD-MEA) System and Recordings

A CMOS-based multi-well HD-MEA system (MaxTwo, MaxWell Biosystems, Zurich, Switzerland) with 26,400 electrodes arranged in a 3.85 × 2.10 mm² sensing area (17.5 µm pitch, 220 × 120 array) was used. Up to 1,020 channels were recorded simultaneously at 10 kHz. The used 24-well system permitted parallel recording of six wells.

All recordings were performed at 37 °C, 5% CO₂ using the MaxLab Live software. Prior to each experiment, a full activity scan of all electrodes was performed (27 configurations, 1 min/configuration). For network recordings, electrodes with the highest average signal amplitude and their surrounding 4×4 grids were selected. For axon scans, 30 high-amplitude electrodes (minimum FR: 0.5 Hz) were selected, complemented by 15 fixed surrounding electrodes to enable reliable spike sorting; the remaining electrodes were recorded sequentially (2 min/configuration).

### Chemogenetic Dose-response Experiments

Medium was exchanged the evening before each dose-response measurement. Controls (non-transduced cultures) were run alongside AAV-transduced cultures, with duplicates of each condition per experiment to obviate any recording order bias. Following a 20 min baseline recording, DCZ was applied in five escalating concentrations (0.01, 0.1, 1, 10, and 100 nм), each preceded by a 10 min equilibration period and followed by a 20 min network recording. Medium was replaced three times after the final recording to ensure a full wash out. Each experiment comprised two time points: weeks 2 and 3 for experiment 1 (n=4 per group; AAV 128-1, AAV 129-1, CTRL) and weeks 1 and 2 for experiment 2 (AAV 128-1, n=3; AAV 129-1, n=3; CTRL, n=2).

### Data Processing and Spike Sorting

All recordings were concatenated per experiment and spike-sorted using SpikeInterface (Buccino et al. 2020) and Kilosort 2.5 (Pachitariu et al. 2024). Units labeled ‘good’ by both Kilosort and the Bombcell quality control pipeline (Fabre et al. 2023) (Table S1) were retained. Single-cell and network features were extracted using the DeePhys toolbox with default parameters (Hornauer et al. 2024). For axon scans, fixed-electrode data were concatenated across scan segments and the dose-response recording before spike sorting, with an additional minimum spike count threshold (50 spikes/segment) for reliable electrophysiological footprint inference.

### Inhibitory Candidate Identification

Putative cell-type labels were assigned, based on FR changes between baseline measurements and the two highest DCZ concentrations. Spike trains were binned into 20-second bins, and a bootstrap permutation test was applied to identify units with a statistically significant increase in firing rate between baseline conditions and DCZ windows. Specifically, for each unit the empirical difference in the mean firing rate (post − pre) was compared against a null distribution of 1,000 permuted differences, obtained by randomly assigning bins to windows of the pooled baseline measurements and post-DCZ spike matrix. Units featuring an empirical difference exceeding the (1 − α/2) percentile of the null distribution were classified as inhibitory candidates, using a significance threshold of α = 10⁻^5^ to minimize false positives.

### Training Label Generation

To build a generalizable classifier, candidate labels derived from chemogenetic activation were translated into a training set in a feature space defined by single-unit electrophysiological properties. For each unit, the autocorrelogram (ACG; 100 ms bidirectional window, 1 ms bins) and the reference waveform (61 samples at 10 kHz, upsampled to 120 kHz using modified Akima cubic Hermite interpolation) were concatenated into a single feature vector. Feature values were z-scored per culture to remove recording-condition variances.

All units obtained through the dose-response recordings were embedded into a 2-dimensional UMAP space using the following hyperparameters: n_neighbors = 50, min_dist = 0.1, spread = 1.0, n_components = 2. The UMAP model was trained using the MATLAB UMAP toolbox (McInnes et al. 2020). The UMAP coordinates of inhibitory candidate units were submitted to an isolation forest (contamination fraction = 0.4, n_estimators = 50) to remove units, the embedding position of which was inconsistent with a coherent inhibitory cluster. The remaining candidates defined the inhibitory training set. To construct the excitatory training set, pairwise Euclidean distances were computed from the inhibitory centroid to all units in the UMAP embedding. Units farther than the 95th percentile of within-candidate distances were considered counterexamples. A subset equal in size to the inhibitory training set was randomly sampled as excitatory training examples. The final training set was thus balanced (1:1 class ratio).

### Cell-Type Classification

The training labels were used to train a supervised UMAP, which optimized the embedding for both geometric structure and class separability. Training data were written to a temporary file and passed to run_umap with the ‘supervised’ option enabled, saving a template for consistent projection of held-out data. Test units (all units not included in the training set) were projected onto the trained embedding, and labels were assigned by the nearest supervisor in the 2-dimensional space. Final cell-type labels were assigned to all units across all cultures.

### E/I Population Activity Decomposition

For each culture, spike trains of units with a confirmed label were binned at 10 ms resolution. The population inhibitory firing rate (*I*) and excitatory firing rate (*E*) were computed as the mean spike counts per bin across all inhibitory and excitatory units. The total population firing rate was the mean value across all labeled units. The instantaneous inhibitory-to-excitatory (*I*/*E*) ratio was computed bin-wise as *I* / *E*; bins with *E* = 0 were excluded from these analyses.

### Classifier Performance Assessment

Classifier performance was evaluated separately for the rat and human *in vitro* datasets. For the rat dataset, accuracy was quantified as the difference between the mean predicted interneuron percentage across single-culture replicates and the corresponding ground truth value obtained from a separate single-cell RNA sequencing experiment. For the human data set, a normalized mean ratio was used to account for systematic prediction offsets, introduced by the pure-culture conditions. Predicted interneuron ratios for each mixed-culture condition were normalized to pure-inhibitory and pure-excitatory culture predictions according to:

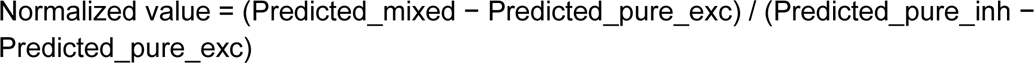

Normalized values were then compared to the expected ground truth ratios, and the resulting differences were averaged across mixed-culture conditions to yield a single performance metric reflecting the mean deviation from the expected interneuron proportion.

### ACG Sub-type Analysis and Classification

To characterize the temporal firing structure of each unit, normalized ACGs were computed over a ±2 second window with 2 ms bins, yielding a symmetric representation of spike-time correlations at positive and negative lags. The zero-lag bin was removed, and the positive-lag half of the ACG was baseline-subtracted by its mean, smoothed with a Gaussian kernel (width = 5 bins), and submitted to peak detection (minimum prominence = 1 SD of the smoothed signal, minimum inter-peak distance = 10 ms).

Oscillatory periodicity was assessed from consecutive inter-peak intervals. A valid frequency estimate required at least three detected peaks, a median inter-peak frequency between 2 and 60 Hz, and a coefficient of variation of consecutive intervals ≤ 0.4. Spectral analysis was performed on the full bilateral (zero-lag–excluded) ACG via FFT, restricted to the 2–60 Hz band. A rhythmicity index (RI) was defined as the ratio of peak spectral power to the mean power across all other frequency bins, and a local spectral SNR was computed as the ratio of peak power to mean power within ± 2 Hz of the peak frequency.

Burst propensity was assessed from the positive-lag ACG prior to mean subtraction. A baseline value was estimated from the terminal 20% of the ACG, and a unit was classified as bursty, when its first detected peak occurred within 100 ms and exceeded three standard deviations of the tail mean value.

Units were assigned to one of four classes according to a decision hierarchy: bursty if the burst criterion was met and the ACG-derived frequency was either undefined or weakly rhythmic (RI < 20); irregular if RI < 20; weakly rhythmic if RI fell between 20 and 50 without a valid periodicity estimate; and oscillatory if all periodicity criteria were satisfied.

### Burst Detection and Alignment

Burst peaks were identified in the smoothened total firing-rate trace using *findpeaks* with minimum peak height and prominence of 0.1 (normalized units) and a minimum inter-peak distance of 6 bins (60 ms). A 3-second window (±150 bins) was extracted around each peak for the total activity, inhibitory activity, and excitatory activity traces. Each burst cutout was aligned to the mean burst shape via circular cross-correlation to correct for sub-bin jitter.

Aligned cutouts were embedded using t-SNE (perplexity = 30) and clustered into two groups by k-medoids partitioning to identify and remove false positives. For cross-culture normalization, the post-peak half of each burst window (bins 101–211 of the 301-bin cutout) was extracted to avoid edge artefacts from alignment, and each culture’s mean burst was normalized by its maximum. The normalized *I*/*E* ratio during the burst was computed analogously and normalized per culture.

### In vivo Data Sets

The rat *in vivo* dataset (Horváth et al. 2021) was recorded using a high-density silicon probe (Fiáth et al. 2018) at a sampling rate of 20 kHz, and subsequently processed exactly like the rat *in vitro* dataset. The mouse *in vivo* Neuropixels dataset (Petersen et al. 2021) was recorded at 30 kHz. This dataset contains 407 optogenetically tagged neurons, including PV+, SST+, excitatory, axo-axonic PV+, and VIP+ cells, whose identity was validated through optogenetic stimulation of Cre-dependent opsin expression in cell-type-specific transgenic mouse lines. Here, preprocessed waveforms and autocorrelograms were already available on the HIPPIE (Gonzalez-Ferrer et al. 2025) github repository (https://github.com/braingeneers/HIPPIE). To ensure compatibility with the *in vitro* dataset, waveforms were upsampled to 120 kHz, and ACG parameters were adjusted to match the corresponding *in vivo* dataset.

### Functional Imaging of Axonal Morphology

Spike sorting results from concatenated axon-scan and dose-response recordings were split post-hoc, with the dose-response data used for cell-type label assignment, and the axon-scan data used for functional morphology reconstruction. To generate electrophysiological footprints (EFs), spike-triggered cutouts were upsampled to 20 kHz, peak-aligned, before computing median spike-triggered averages per configuration and combining them into full-array EFs. Axonal morphology was then inferred from EFs by converting each EF into a spatiotemporal 3D matrix, computing the first derivative to remove slow drifts, applying a spatiotemporal Gaussian filter (sigma: 1), and thresholding by absolute amplitude (< −0.2 μV/ms), relative noise level (< −1.5 × MAD), and maximum propagation velocity (< 1 m/s) from the axon initial segment. The filtered signal was skeletonized using Kimimaro (https://github.com/seung-lab/kimimaro.git) to produce a one-pixel-wide representation of the axonal arbor (scale=1, const=2, anisotropy=(10,10,10), tick_threshold=5, dust_threshold=5). Skeleton branches were validated by computing action potential propagation speed in a sliding window while retaining only branches with velocities of 0.2–0.8 m/s and R² > 0.8; disconnected branches within 175 µm were reconnected if they met this criterion. The full pipeline is publicly available at https://github.com/hornauerp/axon_tracking.

### scRNA-seq Library Preparation, Sequencing and Data Preprocessing

Cryopreserved cells were thawed in a 37 °C water bath and washed gently three times with PBS. Cells were resuspended in 12 µL PBS to obtain a single-cell suspension. Of this volume, 2 µL were used for cell counting, and 10 µL were used as input for scRNA-seq. scRNA-seq were performed using the 10x Genomics’ GEM-X Universal 3’ Gene Expression v4 On-Chip Multiplexing (OCM) platform. GEM generation, library preparation, and sequencing were performed according to the manufacturer’s instructions (10x Genomics User Guide CG000768). Libraries were sequenced on a NextSeq 2000 platform (100-cycle P2), targeting approximately 20,000 reads per cell. Sequencing data were processed using the Cell Ranger v9.0.1 pipeline (10x Genomics) with default parameters. Reads were aligned to the reference genome and transcriptome. For the rat samples, the Ensembl Genome Assembly mRatBN7.2 and corresponding gene annotation file (GTF; release v2.113) were used to generate a custom reference using the Cell Ranger mkref pipeline. FASTQ files were aligned to the reference, and sample demultiplexing as well as cell-by-gene-count matrix generation were performed using the standard Cell Ranger workflow.

### scRNA-seq Data Analysis

To analyze the scRNA-seq data, the cell-by-gene-count matrix generated by the Cell Ranger pipeline was loaded into Seurat (Hao et al. 2024). Cells were filtered based on quality control metrics, retaining cells with a minimum of 400 detected genes. Cells with a high proportion of mitochondrial transcripts (>10%) were also removed. After filtering, 1,564 primary rat cells were retained for downstream analysis. The data were log-normalized using a scaling factor of 10,000. Highly variable genes were identified using the Seurat function FindVariableFeatures (n features = 3,000), after which the data set was scaled. The percentage of mitochondrial gene expression was regressed out during scaling due to its strong influence observed in the initial clustering. Principal component analysis (PCA) was performed using 20 principal components (PCs). The first 10 PCs were used for non-linear dimensionality reduction (UMAP/t-SNE) and subsequent downstream analyses. Cell clustering was performed using the Louvain algorithm with a resolution of 0.5, followed by manual cell type annotation.

Primary rat brain cell types were annotated based on canonical marker gene expression. Neurons were identified by Map2 expression, while neural progenitors and glial cells were marked by Sox2. Glial cells and astrocytes were further supported by expression of Vim, Gfap, and S100b. Dorsal cell fates were marked by Pax6. Glutamatergic neurons were identified by expression of Slc17a6, Slc4a10, Grik2, and Gls, whereas GABAergic neurons were marked by Gad1 and Gad2. Ependymal progenitors were identified by the expression of cilia-associated genes such as Cdhr4 and Ccdc162. Proliferating cells were defined by expression of Mki67 and Top2a.

### Maturation Scoring

To compare the maturation state of primary rodent neurons and human induced neurons, we used published bulk RNA-seq time-course data from human induced glutamatergic neurons at days 10, 21, 28, 35, and 42 after doxycycline induction as a reference (Lin et al. 2025). Neural maturation was quantified based on the collective expression of genes associated with the Gene Ontology (GO) terms Neuron projection, Neurotransmitter secretion, and Synaptic assembly. To calculate the neural maturation score, maturation gene sets were first intersected with genes present in the expression matrix. Expression for the selected genes were then standardized by z-scoring each gene across samples. Finally, the maturation score for each sample was defined as the average z-scored expression of all genes within the module.

### Immunocytochemistry

Immunocytochemical stainings were performed on cover slips that were prepared identically to the HD-MEA cultures, as the 24-well MaxTwo plate is incompatible with conventional microscopes. After aspirating the medium, cells were fixed in 1 mL IC fixation buffer (Invitrogen) for 20 min and washed three times in PBS (Gibco). PBS was then removed, and 1 mL of blocking solution consisting of 10% NDS, 1% BSA, 2% NaN3, and 10% Triton X (all Sigma-Aldrich) in PBS was added to each culture for 30 min. The primary antibody (chicken anti-MAP2, Thermo Fisher Scientific, 1:500) was diluted in a reaction solution consisting of 3% NDS, 1% BSA, 2% NaN_3_, and 10% Triton X in PBS. 250 µL of the primary antibody solution were added to each culture and incubated for 1 hour on a shaker. After washing three times with PBS, the secondary antibody (Alexa fluor 488 anti-chicken, Thermo Fisher Scientific, 1:500) and DAPI (Sigma-Aldrich, 1 µL/mL) diluted in reaction solution were added and incubated on a shaker in the dark for 4 hours. Cultures were again washed three times and kept in PBS. Imaging was done using an upright confocal microscope with a 20x immersion lens (Nikon, Tokyo, Japan). Images were postprocessed using Fiji (Schindelin et al. 2012).

## Acknowledgements

scRNA-seq library preparation and Illumina sequencing were performed at the Genomics Unit at CRG Barcelona.

## Author Contributions (using CRediT taxonomy)

P.H.: Conceptualization (lead), Investigation, Formal Analysis, Writing - Original Draft. L.D.D.: Conceptualization (supporting), Investigation, Formal Analysis, Writing - Review & Editing. H.-C.L.: Funding Acquisition, Investigation, Formal Analysis, Writing - Original Draft. T.G.: Investigation, Formal Analysis, Writing - Review & Editing. M.P.-G.: Investigation, Writing - Review & Editing. M.S.: Funding Acquisition, Conceptualization, Writing - Review & Editing. A.H.: Funding Acquisition, Writing - Review & Editing.

## Data Availability Statement

The classification pipeline will be released as a module of the DeePhys toolbox upon publication (https://github.com/hornauerp/DeePhys). The axon tracking pipeline is publicly available at https://github.com/hornauerp/axon_tracking. Data will be made available by the authors upon reasonable request

