## Supplemental Material for "Functional Cell-Type Identification in Neuronal Networks Using High-Density Microelectrode Arrays"

### Supplementary Material (Tables and Figures)

| Parameter | Value | Parameter | Value |
|---|---|---|---|
| saveSpikes_withoutDuplicates | 1 | maxNTroughs | 1 |
| recomputeDuplicateSpikes | 0 | somatic | 1 |
| detrendWaveform | 1 | minWvDuration | 100 |
| nRawSpikesToExtract | 100 | maxWvDuration | 1500 |
| saveMultipleRaw | 0 | minSpatialDecaySlope | -0.008 |
| decompressData | 0 | minSpatialDecaySlopeExp | 0.01 |
| spikeWidth | 82 | maxSpatialDecaySlopeExp | 0.1 |
| extractRaw | 1 | maxWvBaselineFraction | 0.3 |
| probeType | 1 | firstPeakRatio | 3 |
| computeSpatialDecay | 1 | minTroughToPeakRatio | 0.8 |
| waveformBaselineNoiseWindow | 20 | minWidthFirstPeak | 4 |
| tauR_valuesMin | 1.0000e-03 | minMainPeakToTroughRatio | 5 |
| tauR_valuesStep | 5.0000e-04 | minWidthMainTrough | 5 |
| tauR_valuesMax | 0.003 | isoDmin | 20 |
| tauC | 1.0000e-03 | lratioMax | 0.1 |
| hillOrLlobetMethod | 1 | ssMin | NaN |
| computeTimeChunks | 0 | minAmplitude | 20 |
| deltaTimeChunk | 360 | maxRPVviolations | 0.05 |
| presenceRatioBinSize | 60 | maxPercSpikesMissing | 100 |
| driftBinSize | 60 | minNumSpikes | 18 |
| computeDrift | 0 | maxDrift | 100 |
| waveformBaselineWindowStart | 20 | minPresenceRatio | 0 |
| waveformBaselineWindowStop | 30 | minSNR | 1 |
| minThreshDetectPeaksTroughs | 0.2 | plotDetails | 0 |
| normalizeSpDecay | 1 | plotGlobal | 1 |
| spDecayLinFit | 0 | verbose | 1 |
| ephys_sample_rate | 10000 | reextractRaw | 0 |
| nSyncChannels | 1 | saveAsTSV | 1 |
| gain_to_uV | 6.2 | unitType_for_phy | 1 |
| computeDistanceMetrics | 0 | saveMatFileForGUI | 1 |
| nChannelsIsoDist | 4 | removeDuplicateSpikes | 0 |
| splitGoodAndMua_NonSomatic | 0 | duplicateSpikeWindow_s | 1.0000e-05 |
| maxNPeaks | 2 | | |

**Table S1. Bombcell quality control parameters**

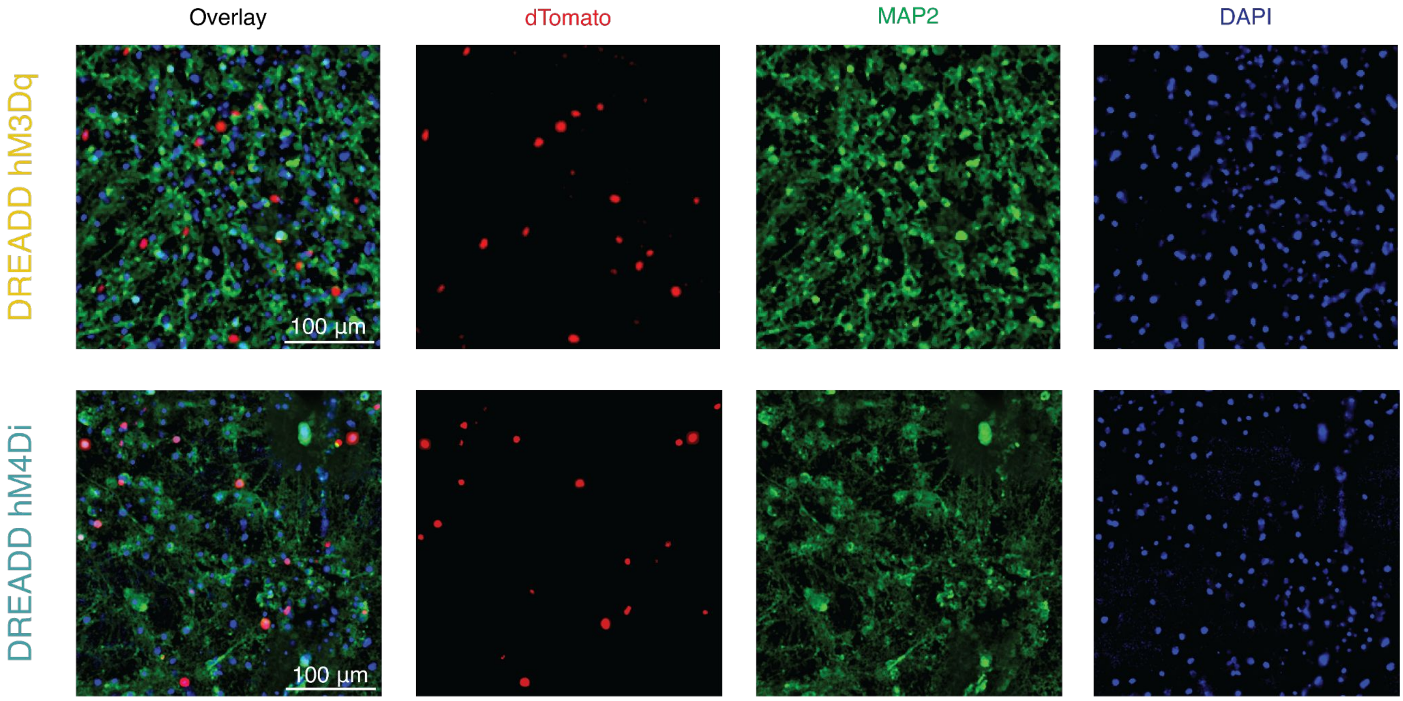


**Figure S1. Immunocytochemical validation of DREADD expression in cortical cultures.** Confocal images of rat cortical cultures transduced with AAV constructs driving interneuron-specific expression of hM3Dq (top row) or hM4Di (bottom row). Columns show, from left to right: overlay, nuclear-localized dTomato reporter (red), MAP2 immunostaining for neurons (green), and DAPI nuclear counterstain (blue). The dTomato signal confirms successful transduction in a subset of neurons, consistent with the interneuron-selective hDlx enhancer driving expression in a minority of the total neuronal population. Scale bars: 100 µm.

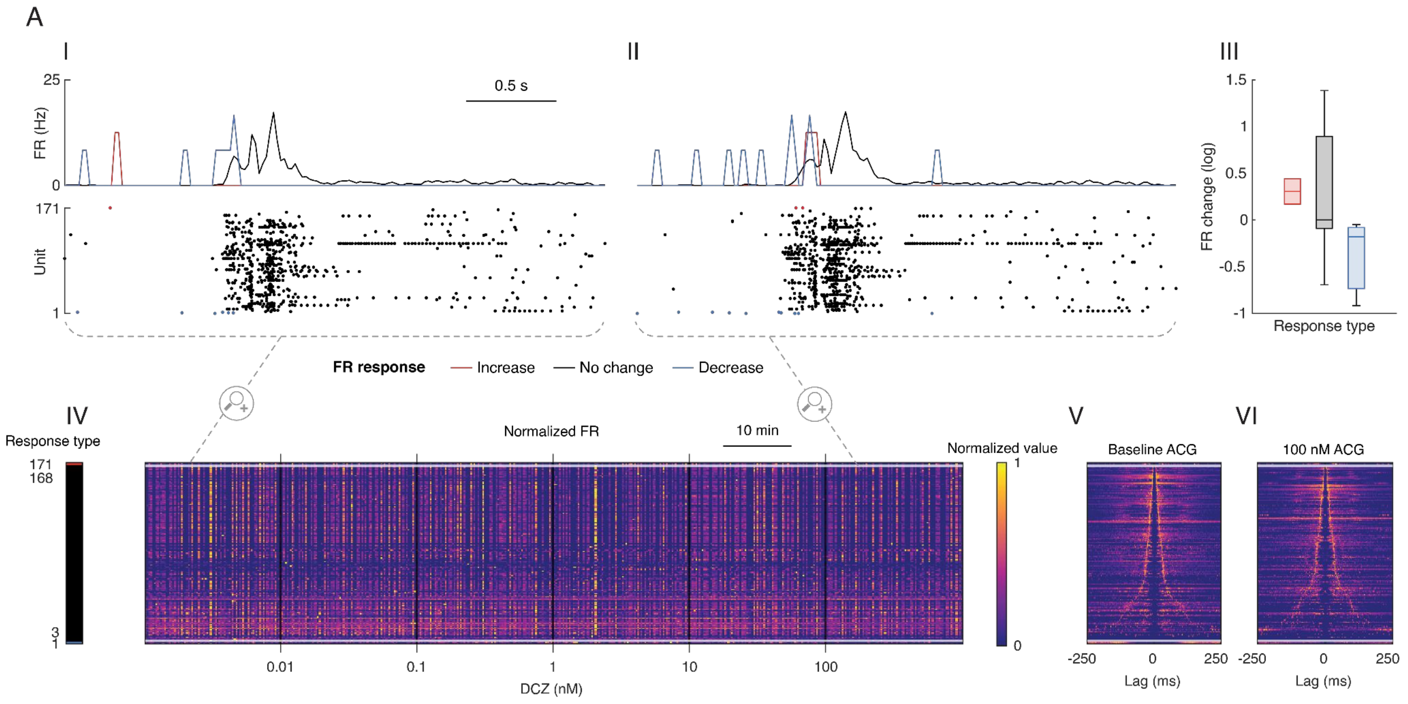


**Figure S2. DCZ application has no effect on non-transduced control cultures.** Analysis of a non-transduced control culture recorded alongside the DREADD-transduced cultures during the DCZ dose-response experiment. **(I)** Representative network burst from the baseline recording. **(II)** Representative network burst at the highest DCZ concentration (100 nM), showing no discernible change in burst structure or firing rate distribution. Units are color-coded by FR response type (Increase: red, No change: black, Decrease: blue). **(III)** Average waveform of all recorded units. **(IV)** Box plots summarizing the FR changes (log) across response types, showing no systematic shift in any subpopulation. **(V)** Heatmap of normalized FR across the full dose-response experiment (15 s bins), confirming stable activity throughout all DCZ concentrations. Units are sorted by response type (left color bar). **(VI–VII)** Autocorrelograms at baseline (VI) and at 100 nM DCZ (VII), showing no change in temporal firing structure (bin size: 1 ms, max lag: 250 ms). Together, these data rule out off-target effects of DCZ on neuronal activity in the absence of DREADD expression.

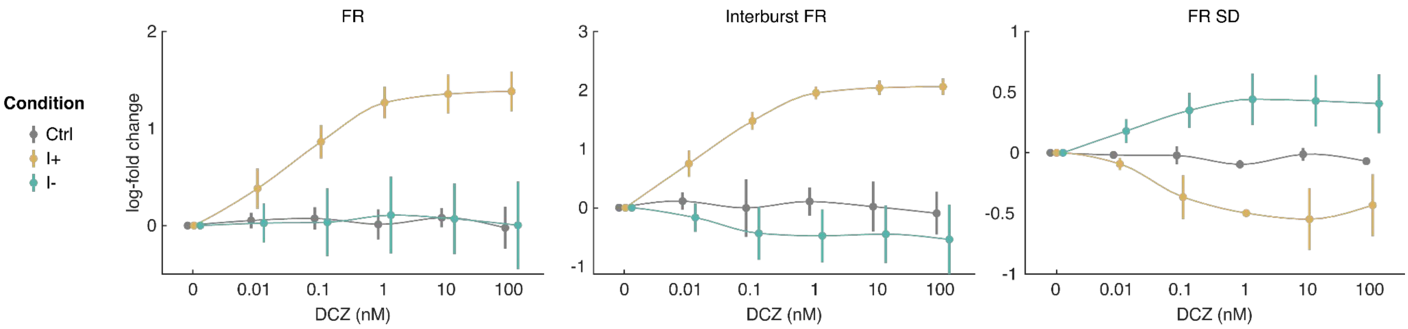


**Figure S3. Dose-response characterization of chemogenetic interneuron modulation.** Dose-response curves quantifying the effect of interneuron stimulation (I+) or inhibition (I−) on culture-wide firing properties compared to non-transduced controls (Ctrl). N = 3 cultures per condition; data are shown as mean log-fold change ± standard deviation (SD), with data points jittered for visual clarity. Statistical significance was assessed using repeated measures analyses of variance (RANOVA) comparing each experimental condition to controls. Left: average firing rate (FR) increased significantly with DCZ dose in the I+ condition (1.4 ± 0.2; $p = 6.03 \times 10^{-5}$) but remained unchanged in the I− condition ($p = 0.954$). Middle: interburst FR showed an even more pronounced increase under I+ (2.06 ± 0.14; $p = 2.38 \times 10^{-5}$), while I− cultures displayed a slight, non-significant decrease (−0.54 ± 0.58; $p = 0.415$), confirming that interneuron activation preferentially increased spiking outside bursts. Right: FR standard deviation decreased under I+ (−0.55 ± 0.25; $p = 0.009$) and increased under I− (0.44 ± 0.21; $p = 8.88 \times 10^{-4}$), reflecting changes in the contrast between intra- and interburst firing states. Control cultures showed no systematic changes across DCZ concentrations for any of the three measures, ruling out off-target effects of DCZ itself.

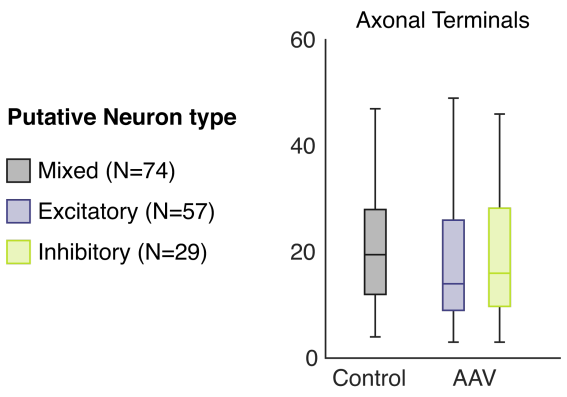


**Figure S4. Axonal terminal count is unaffected by AAV transduction.** Box plots comparing the number of axonal terminals between non-transduced control cultures and AAV-transduced cultures, stratified by putative neuron type: mixed (grey, N = 74), excitatory (blue, N = 57), and inhibitory (yellow, N = 29). No significant differences were observed between conditions ((Kruskal-Wallis test, p=0.979), confirming that DREADD expression does not alter axonal branching complexity.

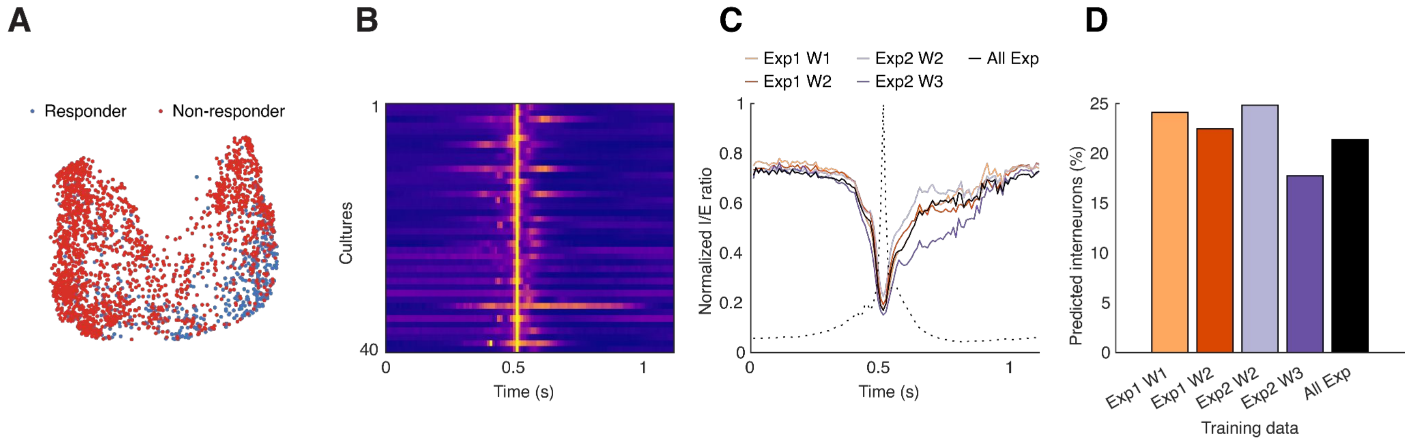


**Figure S5. Predicted interneuron ratio is reproducible across experiments and recording time points. (A)** UMAP embedding of cortical neurons pooled from four chemogenetic stimulation experiments (two biological batches, each recorded at two time points; N = 2,468 units from 14 cultures). Colors indicate responders (red) and non-responders (blue) to DCZ stimulation. **(B)** Heatmap of burst-averaged activity across all cultures from the four experiments. **(C)** Normalized I/E ratio averaged across all cultures and bursts from (B), computed using predicted cell-type labels. Each colored trace corresponds to a classifier trained on a different experiment; the black trace uses pooled training data from all experiments. Dotted line indicates the corresponding average network activity. **(D)** Predicted interneuron percentage when using each individual experiment as training data. The predicted fraction is comparable across experiments (range: ~18–27%), with the pooled classifier (black) yielding ~20%. N = 7,333 units pooled across all experiments and conditions. Exp: experiment (biological batch); W: week of recording.

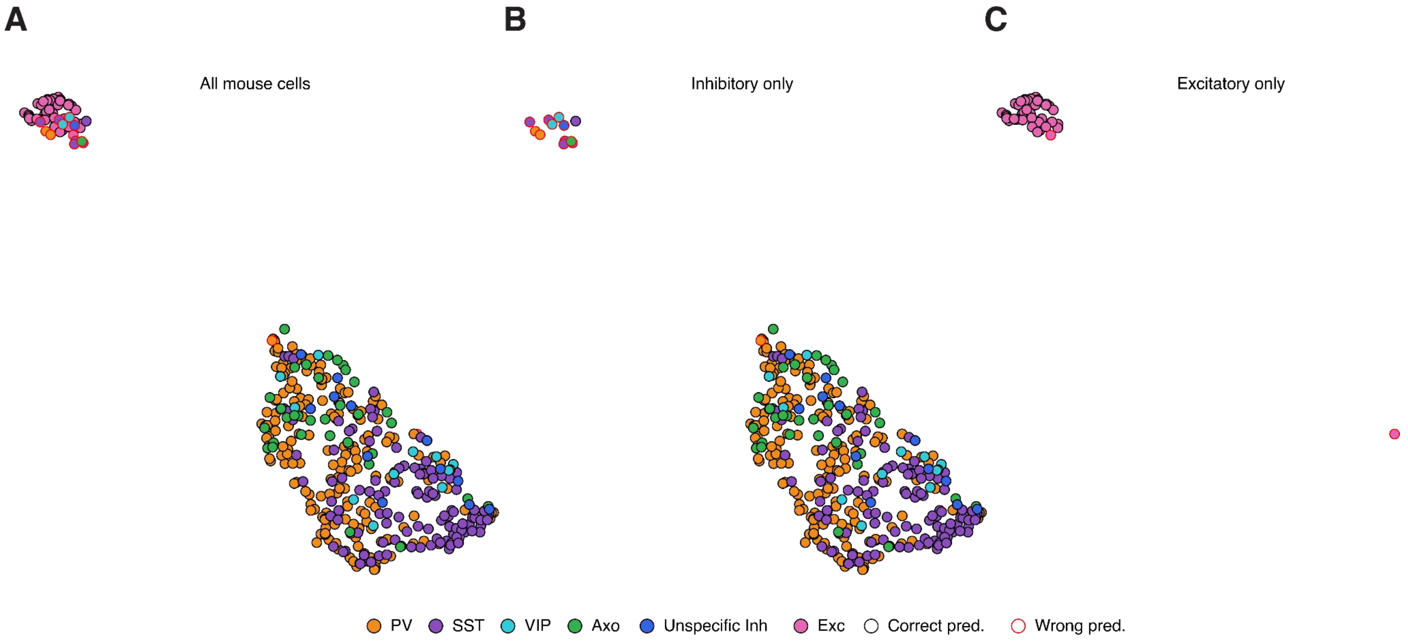


**Figure S6. Classifier performance on the mouse *in vivo* Neuropixels dataset.** The classifier trained on the *in vitro* rat dataset was applied without retraining to the mouse *in vivo* dataset. **(A)** Supervised UMAP embedding of all units, colored by optogenetic ground-truth label: excitatory (Exc, pink) and inhibitory subtypes (PV, orange; SST, purple; VIP, cyan; Axo, green; Unspecific Inh, dark blue). Black outlines indicate correct predictions; red outlines indicate misclassifications. **(B)** Inhibitory neurons only. **(C)** Excitatory neurons only. Inhibitory subtypes predominantly cluster together in the lower portion of the embedding, while excitatory neurons occupy the upper left. Notably, a small number of inhibitory neurons embed within the predominantly excitatory cluster, creating a region of overlap where excitatory neurons are more likely to be misclassified. This asymmetry likely reflects the training set composition, in which inhibitory labels were derived from the chemogenetic experiment while excitatory labels were generated by exclusion, resulting in higher confidence for inhibitory than excitatory assignments.

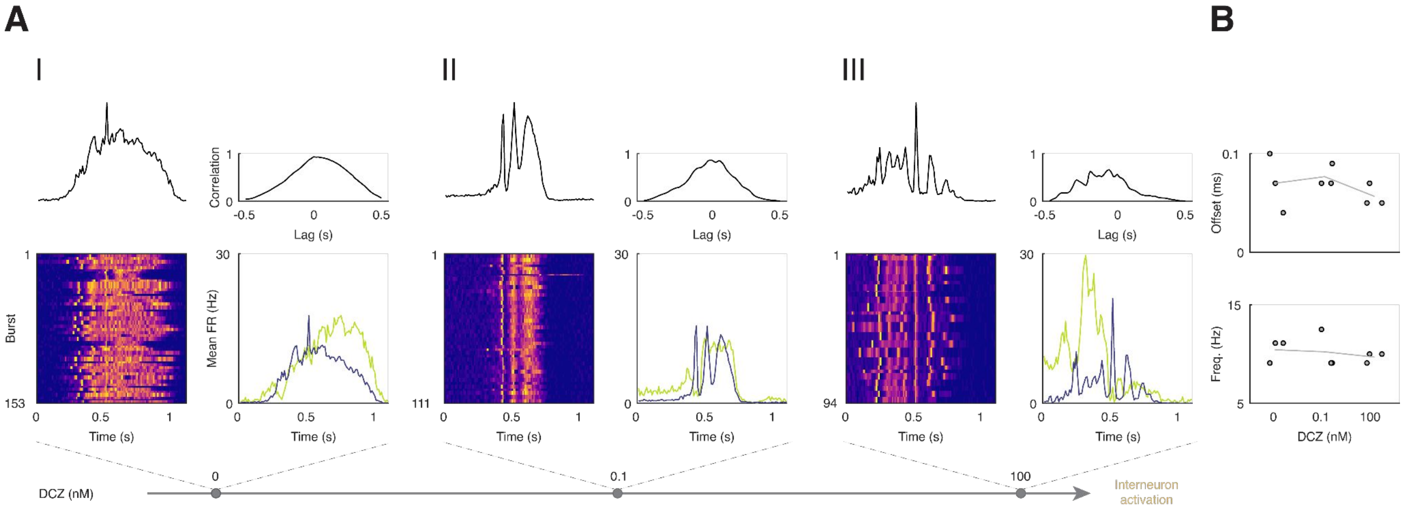


**Figure S7. Burst dynamics in immature cortical cultures are more strongly affected by interneuron activation. (A)** Burst analysis as described in Figure 4A, applied to cortical cultures at DIV 10 (N = 3). Panels **(I)**, **(II)**, and **(III)** correspond to baseline, 0.1 nM DCZ, and 100 nM DCZ recordings, each showing the I/E cross-correlogram (top), burst-aligned activity heatmap (bottom left), and mean excitatory (blue) and inhibitory (yellow) firing rate traces (bottom right) of a representative culture. Compared to mature cultures (Figure 4B), immature networks show more pronounced restructuring of burst dynamics with increasing interneuron activation. **(B)** Summary of offset (top) and firing frequency (1/period, bottom) across DCZ concentrations.

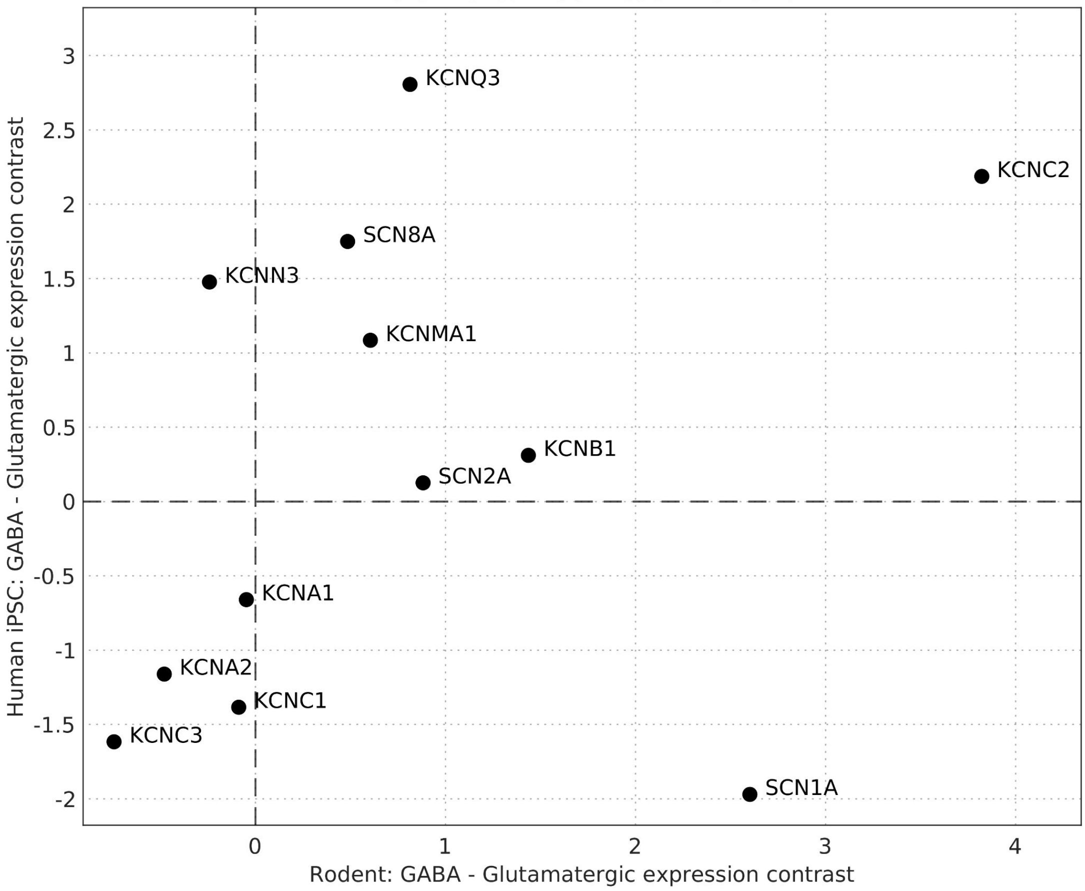


**Figure S8. scRNAseq differential expression levels of genes involved into AP shape explanation between rodent and human neurons.** Interestingly, SCN1A shows lower expression in human GABAergic than glutamatergic cells (-1.971) while higher expression in rodent GABAergic than glutamatergic cells (2.602). Conversely, KCNN3 shows higher expression levels in human GABAergic than glutamatergic cells (1.477) while lower expression in rodent GABAergic than glutamatergic cells (-0.241). Expression contrast was calculated as the difference in variance-stabilized gene expression between GABAergic and glutamatergic neuronal populations.
